# Biochemical and structural characterization of the fused *Bacteroides fragilis* NFeoAB domain reveals a role for FeoA

**DOI:** 10.1101/2021.09.29.462438

**Authors:** Alex E. Sestok, Janae B. Brown, Juliet O. Obi, Sean M. O’Sullivan, Elsa D. Garcin, Daniel J. Deredge, Aaron T. Smith

## Abstract

Iron is an essential element for nearly all organisms, and under anoxic and/or reducing conditions, Fe^2+^ is the dominant form of iron available to bacteria. The ferrous iron transport (Feo) system has been identified as the primary prokaryotic Fe^2+^ import machinery, and two proteins (FeoA and FeoB) are conserved across most bacterial species. However, how FeoA and FeoB function relative to one another remained enigmatic. In this work we explored the distribution of *feoAB* operons predicted to encode for a fusion of FeoA tethered to the soluble N-terminal, G-protein domain of FeoB via a connecting linker region. We hypothesized that this fusion might poise FeoA to interact with FeoB in order to affect function. To test this hypothesis, we cloned, expressed, purified, and biochemically characterized the soluble NFeoAB fusion protein from *Bacteroides fragilis*, a commensal organism implicated in drug-resistant peritoneal infections. Using X-ray crystallography, we determined to 1.50 Å resolution the structure of *Bf*FeoA, which adopts an SH3-like fold implicated in protein-protein interactions. In combination with structural modeling, small-angle X-ray scattering, and hydrogen-deuterium exchange mass spectrometry, we show that FeoA and NFeoB indeed interact in a nucleotide-dependent manner, and we have mapped the protein-protein interaction interface. Finally, using GTP hydrolysis assays, we demonstrate that *Bf*NFeoAB exhibits one of the slowest known rates of Feo-mediated GTP hydrolysis and is not potassium-stimulated, indicating that FeoA-NFeoB interactions may function to stabilize the GTP-bound form of FeoB. Our work thus reveals a role for FeoA function in the fused FeoAB systems and suggests a broader role for FeoA function amongst prokaryotes.

## INTRODUCTION

Nearly all living organisms rely on iron acquisition and utilization for vital cellular processes from aerobic cellular respiration, N_2_ fixation, gene regulation, and DNA biosynthesis (1-3). Given the versatile functionality of iron, this element may be used as an ionic cofactor and bound by biological macromolecules such as in ribonucleotide reductases (4), utilized in [Fe-S] clusters such as those in nitrogenase (5) or the electron transport chain (6), and even chelated in protoporphyrin-IX (heme) and bound to O_2_-carrying proteins such as hemoglobin and myoglobin (7). However, a prerequisite of iron incorporation into proteins is the acquisition of this element, which can be challenging. While ferric iron (Fe^3+^) is predominantly present in oxic environments, it is highly insoluble (K_sp_ *ca*. 10^−18^ M at pH 7.0) (1,8). Conversely ferrous iron (Fe^2+^) is much more soluble than Fe^3+^ (K_sp_ up to 0.1 M at pH 7.0), but ferrous iron is readily susceptible to oxidation and may be incredibly toxic to the cell via Fenton-like chemistry, if unregulated (1,8,9). As a result, organisms must exert both high energy and tight control over the iron acquisition process.

Historically, much work has been done to elucidate bacterial mechanisms of ferric iron and heme transport due to the link of these iron acquisition processes to pathogenesis. For example, it is well established that bacteria secrete small molecules called siderophores into the extracellular space to acquire ferric iron. These molecules have a high affinity for Fe^3+^ (K_aff_ ≥ 10^30^ M^-1^) and allow bacteria to compete against host Fe^3+^-binding proteins for ferric iron (8,10,11). Once acquired and delivered into the cytoplasm, Fe^3+^ can be released by degrading the siderophore or through reducing Fe^3+^ to Fe^2+^, which is accomplished by ferric iron reductases (8,10-12). It is also well known that bacteria employ dedicated transport systems to acquire heme. Heme acquisition is achieved through the use of hemophores, proteins that bind heme specifically and allow bacteria to compete for heme with host heme-binding proteins (13-15). Once delivered into the cytoplasm via a number of membrane-imbedded transporters, heme oxygenases then degrade heme to release iron for incorporation into proteins and metabolic enzymes (13-15).

In addition to Fe^3+^ and heme, bacteria can also transport and utilize Fe^2+^, although this process is far less well understood. The most widespread, dedicated prokaryotic machinery for Fe^2+^ import is the ferrous iron transport (Feo) system. The Feo system was first identified in 1987, and while *E. coli* has a “canonical” arrangement of three genes (*feoA*/*B*/*C*), an arrangement in which only the *feoA* and *feoB* genes are present is far more common in bacteria (16-18). The function of FeoA is unknown, but we do know that FeoA is an ≈8 kDa, cytoplasmic β-barrel protein comprising an Src homology 3 (SH3)-like fold, which is commonly involved in protein-protein interactions. Given its structure, FeoA has been hypothesized to interact with FeoB to affect function (2,3,19,20). FeoB is an ≈85 kDa transmembrane protein consisting of a G-protein domain, a guanine dissociation inhibitor (GDI) domain, and a transmembrane (TM) domain. The G-protein domain is responsible for binding and hydrolyzing GTP (21), though recent studies have shown that some FeoB proteins are also capable of hydrolyzing ATP (22,23). The GDI domain links the G-protein domain to the TM region and has been shown to increase the binding affinity of GDP (24). Together, the G-protein domain and the GDI domain comprise what is termed NFeoB. The TM region has not been structurally characterized but is likely the domain through which Fe^2+^ is translocated (25-27).

Despite our lack of mechanistic information, several studies have demonstrated the importance of the Feo system for the intracellular colonization, survival, and virulence of many pathogens. These infectious bacteria include, but are not limited to, *Legionella pneumophila* (28), *Campylobacter jejuni* (29), *Francisella tularensis* (30), avian pathogenic *Escherichia coli* (31), *Shigella flexneri* (32), and *Streptococcus suis* (33). Interestingly, in some pathogens such as *Porphyromonas gingivalis* (the causative agent of gingivitis) (34) and *Bacteroides fragilis* (a commensal organism implicated in drug-resistant peritoneal infections) (35,36), the *feo* operon is predicted to encode a single FeoAB fusion protein in which FeoA is naturally tethered to the soluble G-protein domain of FeoB (2,3). The presence of these fusion proteins in bacterial genomes strongly suggests that FeoA and FeoB are meant to interact and to work in concert with one another. However, these fusions had yet to be studied at the protein level, representing a clear opportunity to probe uniquely into Feo structure and function.

Herein, we provide the first biochemical and biophysical characterization of the soluble domain of the *Bacteroides fragilis* FeoAB fusion protein (*Bf*NFeoAB). Using genomic data, we demonstrate that FeoAB fusion proteins are more widespread than initially thought, and that these fusions appear to be predominantly found in host-associated bacteria. We subsequently cloned, expressed, and purified *Bf*NFeoAB for X-ray crystallography, small-angle X-ray scattering, and hydrogen-deuterium exchange mass spectrometry. Using these biophysical approaches, we show that *Bf*FeoA bears a conserved SH3-like fold, that apo *Bf*NFeoAB adopts an open, extended conformation in solution, and that interactions of *Bf*FeoA with *Bf*NFeoB occur in a nucleotide-mediated fashion that occludes essential parts of the G-protein domain. Lastly, we use nuclear magnetic resonance spectroscopy to show that the FeoA-NFeoB fusion exhibits exceedingly slow rates of GTP hydrolysis and is not potassium stimulated. Combined, these data suggest a mechanism in which FeoA interacts with NFeoB in a nucleotide-mediated manner, and we hypothesize this function is to attenuate GTP hydrolysis.

## RESULTS

### Distribution of FeoAB fusion proteins

Queried nearly a decade ago, a previous study estimated (based on only 33 sequenced bacterial genomes) that ≈3% of all *feo* operons encode for a fusion of the FeoA protein to the N-terminal, soluble G-protein domain of NFeoB (2,19). To update the prevalence and distribution of the fusion proteins across bacteria, we leveraged more extensively sequenced genomes and utilized the InterPro Database to search for predicted protein architectures containing FeoA (IPR007167). Consistent with the notion that FeoA commonly functions as a single, stand-alone polypeptide, ≈88% of the *feoA* open reading frames (ORFs) appear to be discontinuous of the *feoB* ORF (25,203 of 28,444 sequences). Interestingly, the encoded FeoA protein is predicted to be fused to another FeoA protein, (*i*.*e*., FeoA-FeoA or FeoA-FeoA-FeoA) in ≈3% (791 of 28,444) and ≈0.1% (29 of 28,444) of gene architectures, respectively. The remaining 2,421 genes (*ca*. 8.5% of the sequenced bacterial genomes) are a single, continuous *feoAB* ORF, predicted to encode a single polypeptide in which FeoA is fused to the N-terminal G-protein of FeoB, indicating a higher prevalence of this arrangement than initially thought (Fig. 1). Though there is some diversity in the predicted gene architectures amongst these fusion proteins, a majority (≈92%; 2,225 sequences) have four predicted domains in total: FeoA, the G-protein domain, the GDI domain, and TM region (Fig. 1A). Very few (<3%; 66 sequences) lack a GDI domain. Even more rare, 44 sequences are composed of solely FeoA and the G-protein domain of FeoB, while 31 sequences are composed solely of FeoA and an intact NFeoB (*i*.*e*., no transmembrane region; Fig. 1A). However, given the poor conservation of these truncated sequences and their likely lack of function, it is possible that these may be sequencing errors.

**Fig. 1.**
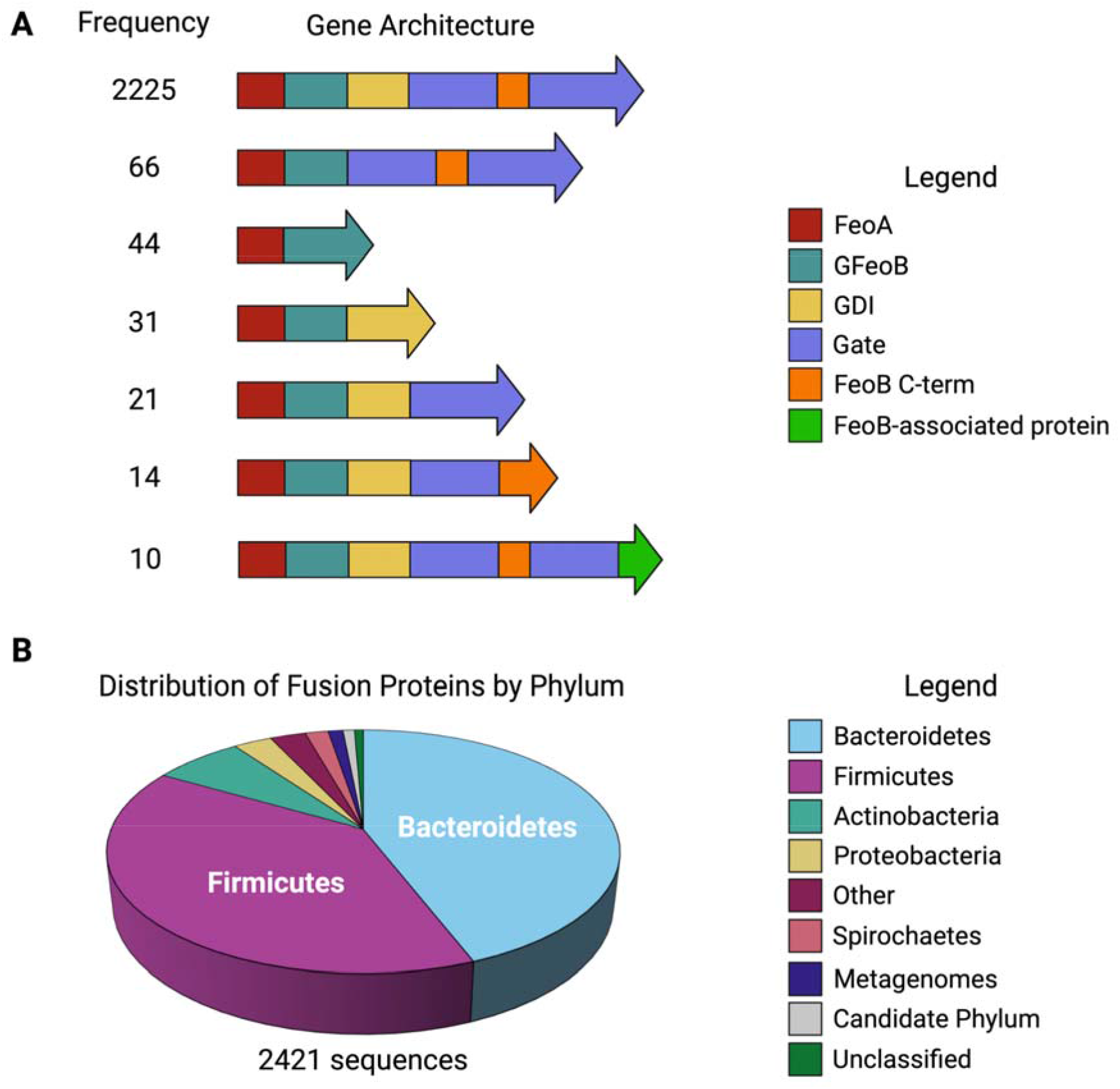
Organization and distribution of FeoAB fusion proteins. **A**. Gene architectures of FeoAB fusion proteins in the InterPro Database as of Feb. 2021. Most FeoAB fusion proteins are predicted to be composed of four domains: FeoA (red), the G-protein domain (teal), the GDI domain (yellow), and the transmembrane region, which comprises the gate domain (purple) and a C-term extension (orange). Very few FeoAB fusion proteins lack the transmembrane region, and these predicted proteins could represent sequencing errors. Though rare, 10 FeoAB fusion proteins are predicted to be fused to an FeoB-associated Cys-rich membrane protein of unknown function (green). **B**. FeoAB fusion proteins are predominantly distributed in the *Bacteroidetes* (light blue) and *Firmicutes* (purple) phyla. Even fewer are present in *Actinobacteria* (teal), *Proteobacteria* (gold), and *Spirochaetes* (salmon). FeoAB fusion proteins have been discovered in metagenomes (dark blue), uncultured bacteria (gray), and unclassified bacteria (green). Other (dark pink) refers to bacterial phyla with fewer than 10 discovered FeoAB fusions. Figure created with BioRender.

We then analyzed the organismal distribution of these fusion sequences to probe their distribution across bacterial phyla (Fig. 1B). FeoAB fusions appear to be predominantly distributed in the *Bacteroidetes* phylum (≈44%) and the *Firmicutes* phylum (≈39%), which are Gram-negative and Gram-positive bacteria, respectively. These organisms constitute a large portion of the human gut microbiome where they facilitate the breakdown of polysaccharides, such as cellulose and starches, and provide the host with a substantial energy source (37-39). Even fewer sequences are found in *Actinobacteria* (≈7%), *Proteobacteria* (≈3%) and *Spirochaetes* (≈2%), members of which are also found in the human gut microbiome (37,38). As several of these organisms live in acidic and/or anoxic conditions, it is possible that these bacteria leverage unique properties of the FeoAB fusion proteins to provide a major portion of the organism’s iron stock. Moreover, as it has been suggested that FeoA and FeoB likely interact with one another at the NFeoB domain, we were then motivated to characterize an NFeoAB fusion at the protein level for the first time.

### Expression and purification of BfNFeoAB

As these FeoAB fusion proteins have not been characterized *in vitro*, we sought to clone, express, purify, and to characterize an NFeoAB fusion to gain insight into FeoA function. Of the >2,000 sequences available, we chose *Bacteroides fragilis* (a representative of the *Bacteroidetes* phylum), which is a commensal, anaerobic, non-spore forming bacterium that colonizes the human gut. In order to investigate the structure and function of one of these N-terminal fusions, the codon-optimized gene corresponding to the N-terminal soluble domain of *Bf*FeoAB (*Bf*NFeoAB; amino acid residues 1-438) (Fig. 2A) was subcloned into a pET-based plasmid and expressed heterologously in *E. coli* with a C-terminal (His)_6_ tag for ease of purification. After sonication and lysate clarification, the soluble *Bf*NFeoAB was initially purified via immobilized metal affinity chromatography. After just one round of column chromatography, large quantities of significantly pure protein (≈80-100 mg/L culture) could be obtained. We then assessed the homogeneity of *Bf*NFeoAB by subsequent size-exclusion chromatography (SEC) on Superdex 200 (Fig. 2B). Interestingly, while a small portion of *Bf*NFeoAB migrated as an apparent trimeric species (estimated <10%; Fig. 2B), the vast majority of the protein (estimated >90%) was monomeric under these conditions (Fig. 2B). This observation contrasts with some theories that NFeoB exists as only a trimeric species (40,41). Concentration of the monomeric species and subsequent reinjection preserved the monomeric oligomer, indicating that a dynamic equilibrium was not operative on this time scale (several hours; data not shown). We then assessed the purity of our SEC-purified *Bf*NFeoAB by SDS-PAGE, which migrated similarly to the estimated MW (≈53 kDa; Fig. 2C), consistent with our SEC data. This monomeric protein was estimated to be >95% pure and was used for all subsequent biochemical and biophysical analyses.

**Fig. 2.**
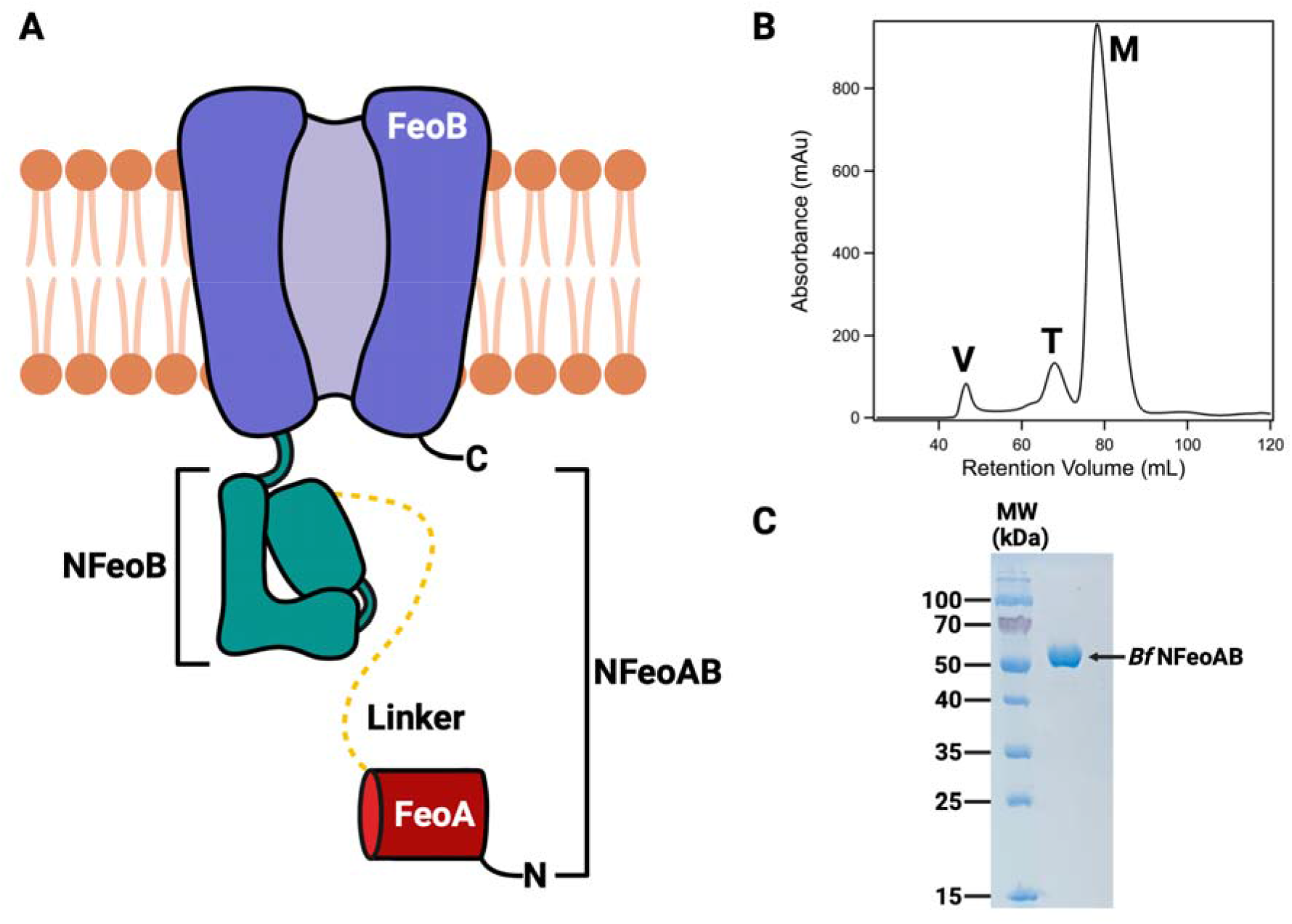
Predicted domain topology of the *Bf*NFeoAB construct used in this work and its purification. **A**. Cartoon representation of the FeoAB fusion protein from *Bacteroides fragilis*. The FeoA protein (red) is covalently tethered to NFeoB (teal) through the G-protein domain by a predicted 44-amino acid linker region (dashed yellow line). The soluble NFeoAB domain is tethered to the transmembrane region of FeoB (purple). Labels ‘N’ and ‘C’ refer to the N- and C-termini, respectively. **B**. SEC purification of *Bf*NFeoAB on a 120 mL Superdex 200 column. The majority of *Bf*NFeoAB is monomeric (≈80 mL retention volume; ‘M’), while no more than 10% of *Bf*NFeoAB is either trimeric (≈70 mL retention volume; ‘T’) or aggregated (≈45 mL retention volume; ‘V’). **C**. Based on SDS-PAGE, *Bf*NFeoAB is estimated to be >95% pure after SEC in panel **B**. Figure created with BioRender.

### Crystallization of BfFeoA

Given the high purity and homogeneity of *Bf*NFeoAB, we next sought to crystallize this protein in the apo and nucleotide-bound forms. Despite exhaustive initial crystallization trials in the presence and absence of nucleotides, drops in sparse matrix screens remained mostly clear, even after testing protein concentration, protein state (± His tag), nucleotide type, and even nucleotide composition. However, after ≈11 months of equilibration, crystals were obtained in ammonium sulfate and di-potassium phosphate that were further optimized by grid screening. After multiple months, medium-sized rectangular crystals appeared that were looped, cryoprotected, and screened for diffraction.

Despite the age of the crystals, diffraction was routinely observed to <2 Å resolution, with our best datasets extending to 1.50 Å (Table S1). After data processing, we initially attempted to phase our data using molecular replacement (MR) of established NFeoB models. However, given the small monoclinic unit cell (*C*_121_; a=92.58 Å; b=29.55 Å, c=67.43 Å; α=90°; β=128.89°; and γ=90°; Table S1), it was clear that the intact fusion protein could not be present within the lattice with a reasonable solvent content. Under this assumption, we were then able to phase our data by MR using *Clostridium thermocellum* FeoA (PDB ID 2K5L) as an input search model. Initial refinement revealed the presence of two molecules of only the FeoA domain (*Bf*FeoA) in the asymmetric unit (ASU). After iterative rounds of rebuilding and refinement, our *Bf*FeoA model converged with *R*_w_ = 0.192 and *R*_f_ = 0.236 (Table S1).

Our crystal structure of *Bf*FeoA comprises residues 1-74 of the *Bf*NFeoAB polypeptide (PDB ID 7R7B). As visible in Fig. 3A, *Bf*FeoA adopts the β-barrel, SRC Homology 3 (SH3-like) fold that has been observed for other FeoA proteins (19,20,42). The β-barrel is composed of five β-strands, while two α-helices make up the clamp region of *Bf*FeoA, which comprises a series of hydrophobic residues (Phe^23^, Ile^27^, Met^30^, Ile^59^, and Leu^61^) that we have hypothesized to be important for mediating FeoA-NFeoB interactions (Fig. 3B) (20). An additional 3_10_-helix at the beginning of the N-terminus contacts both the final α-helix and β-strand that feed out of and into the hydrophobic clamp, respectively. Residues comprising both the linker region between FeoA and NFeoB (Fig. 1A) are completely absent from the structure, likely a result of proteolytic degradation in the crystallization drop over time. While sequence analysis of the FeoA polypeptide does not suggest this is a common proteolytic site, the new C-terminus created after proteolysis is unambiguously present in the electron density (Fig. S1). Although not what we initially set out to crystallize, this structure nevertheless represents the first structure of a portion of the *B. fragilis* Feo system.

**Fig. 3.**
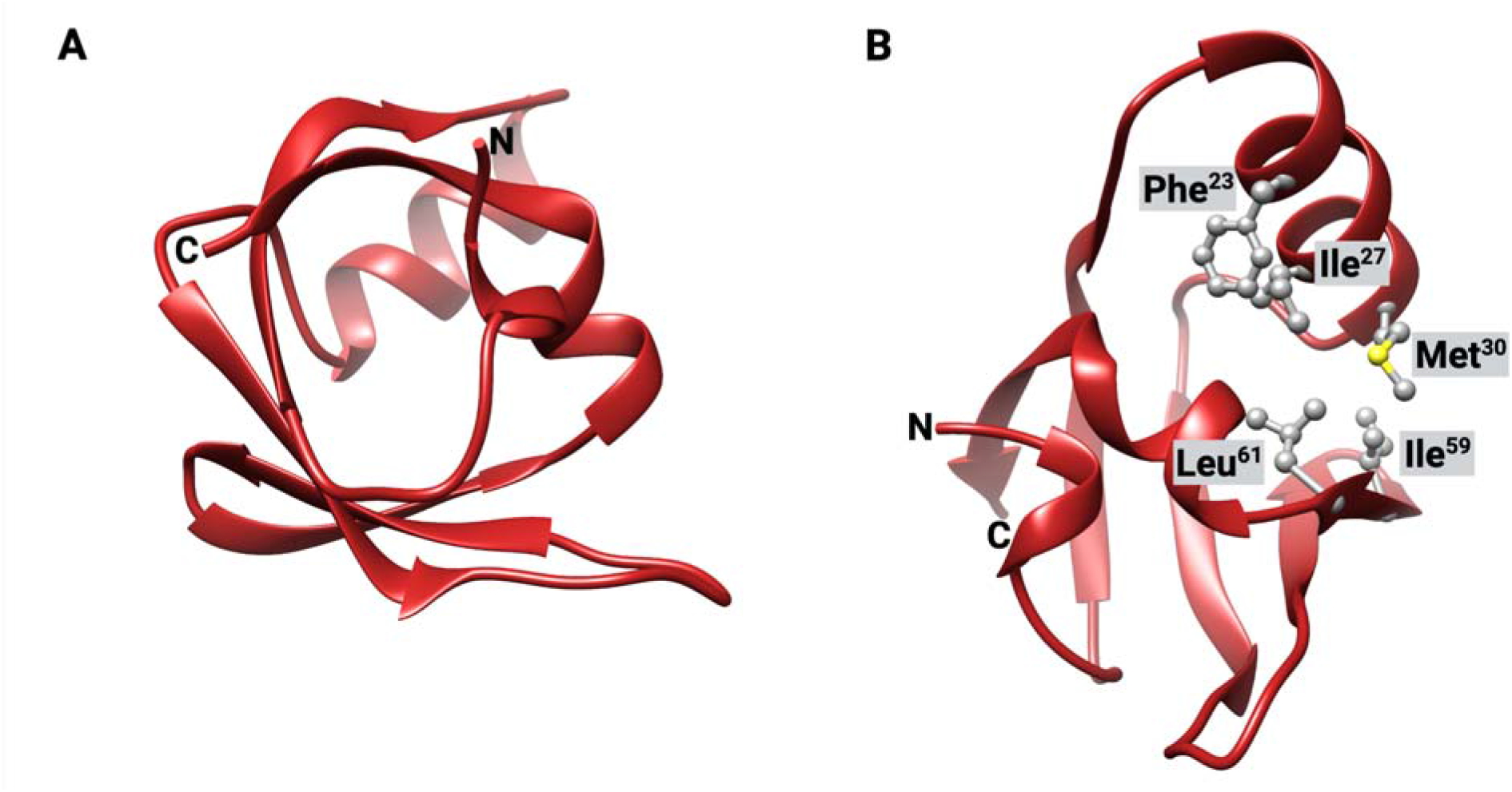
Crystal structure of *Bf*FeoA. **A**. Similar to other FeoA proteins structurally characterized, *Bf*FeoA adopts an SH3-like fold, which is characterized by a small β-barrel. **B**. Like observed in the structure of *Kp*FeoA (PDB ID 6E55), *Bf*FeoA contains a “C-shaped” clamp region lined with hydrophobic residues (shown in ball and stick), composed of Phe^23^, Ile^27^, Met^30^, Ile^59^, and Leu^61^. This view represents a 100° rotation about the y-axis and a 40° rotation about the x-axis of panel **A**. Labels ‘N’ and ‘C’ refer to the N- and C-termini, respectively. Figure created with BioRender.

### Homology Modeling

Since our crystallization screening only yielded FeoA crystals, we instead turned to homology modeling to predict the structure of the intact, soluble NFeoAB fusion protein from *B. fragilis*. Using homology approaches applied by the Robetta server (43,44), five models were generated with high confidence. There appeared to be very little difference amongst the predicted three-dimensional folds of each model, with the exception of the placement of FeoA and the intervening linker region vis-à-vis NFeoB. A representative model is shown in Fig. 4A, which is most consistent with our in solution biophysical data (*vide infra*).

**Fig. 4.**
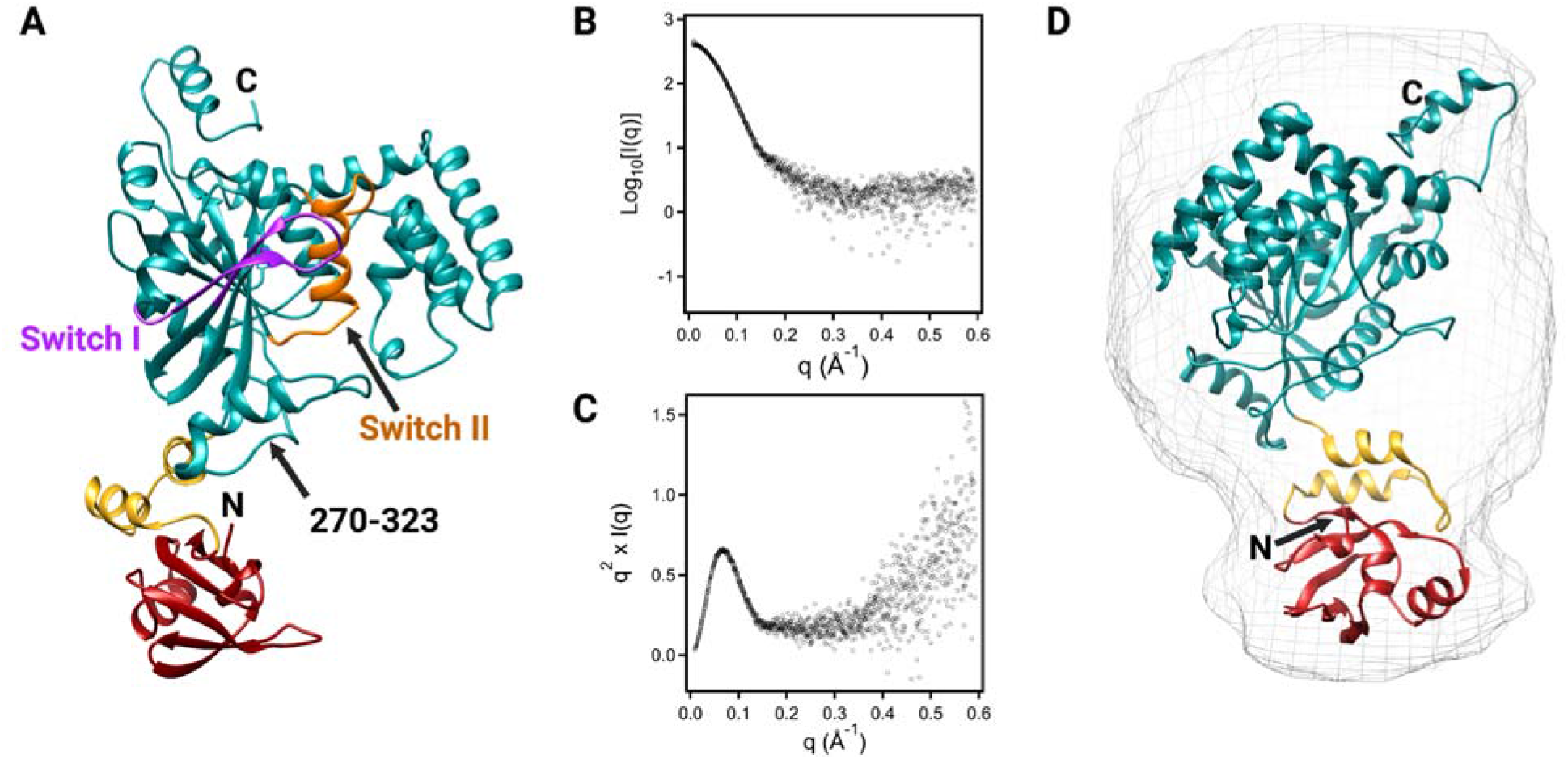
Modeling and SAXS data of apo *Bf*NFeoAB. **A**. Representative Robetta model of apo *Bf*NFeoAB. FeoA (red) is covalently tethered to NFeoB (teal) through a flexible linker region (yellow). Similar to other NFeoB proteins, the Robetta model reveals that *Bf*NFeoAB has both a Switch I region (purple) and a Switch II region (orange). Residues 270-323 represent an additional α-helix and unstructured loop not present in other NFeoBs. **B**. The log_10_ plot of apo *Bf*NFeoAB SEC-SAXS data indicates that the protein is monodisperse with negligible aggregation. **C**. The Kratky plot derived from SEC-SAXS data of apo *Bf*NFeoAB is bell-shaped, indicating a well-folded protein. As the curve does not return to 0 at high q values, these data indicate flexibility within the protein. **D**. Overlay of our best-fit Robetta model of apo *Bf*NFeoAB with the *ab initio* envelope (gray mesh) generated from the SEC-SAXS data. The envelope is elongated, and FeoA points away from the G-protein and GDI domains in the apo form. Labels ‘N’ and ‘C’ refer to the N- and C-termini, respectively. Figure created with Biorender.

We then compared our homology models of *Bf*NFeoAB to structurally characterized Feo proteins to determine the locations of each domain of *Bf*NFeoAB. As our newly determined FeoA structure was not part of the homology modeling, we first superposed the FeoA portion of our Robetta model (1-74) with our *Bf*FeoA crystal structure to determine how similar the structures are. Both structures aligned well with a root-mean-square deviation (R.M.S.D.) of 0.786 Å over 67 C_α_s that comprise the core SH3-like fold. The largest deviations were found in two loop regions spanning amino acids 18-23 and 42-50. We then aligned our models with apo *Ec*NFeoB (PDB ID 3I8S, chain A) and were able to define the *Bf*NFeoB region as spanning residues 110-447, while the linker region spans residues 75-109 (Fig. 4A). Interestingly, the linker region was modeled as two α-helices connected by a short, unstructured loop in all Robetta-generated models (Fig. 4A), suggesting more structure in the linker than we initially suspected.

We then determined the locations of key G-protein motifs in our *Bf*NFeoAB models using multiple sequence alignments (Fig. S2) and structural comparisons to apo *Ec*NFeoB. Based on these comparisons, we determined that the G1 motif is located at positions 118-125 (GNPNCGKT), the G2 motif is located at position 145 (T), the G3 motif is located at positions 164-167 (DLPG), the G4 motif is located at positions 224-227 (NMYD), and the G5 motif is located at positions 254-259 (CKRNIG). *Bf*NFeoAB also contains a PxxP sequence in the G-protein domain, similar to other FeoB proteins. In *E. coli*, this sequence is located at positions 144-147, which corresponds to positions 248-251 in *Bf*NFeoAB. Additionally, we predict the Switch I region (Fig. 4A, purple) to be located at position 134-150 and the Switch II region (Fig. 4A, orange) to be located at positions 170-191. Structural superposition, this time with that of the *Bf*NFeoB models and apo *Ec*NFeoB (PDB ID 3I8S, chain A), resulted in an R.M.S.D. of 0.966 Å over 170 C_α_s in the core of the G-protein, although notable exceptions are observed in key dynamic regions. The largest deviations were observed from residues 133-146 (the Switch I region), residues 154-159 (between the G2 and G3 motifs), the G5 motif, and in residues 270-323 that are modeled as an additional α-helix and an unstructured loop between the G-protein and the GDI domain in the *Bf*NFeoAB models but are not present in the apo *Ec*NFeoB structure. Thus, our full-length Robetta models of apo *Bf*NFeoAB are in good agreement with structural data on native and homologous Feo proteins.

### Small-Angle X-Ray Scattering

Since we were unable to crystallize intact *Bf*NFeoAB in the presence or absence of nucleotides, we instead used small-angle X-ray scattering (SAXS) to determine its solution structure, and to compare the experimental solution structure to our Robetta models. Our initial high-throughput (HT) SAXS experiments on apo *Bf*NFeoAB (Fig. S3A) revealed homogenous protein with minimal aggregation in protein samples at low concentrations (Fig. S3B), and suggested an elongated conformation (Fig. S3C), similar to our homology models. This observation was confirmed for apo *Bf*NFeoAB by using SEC-SAXS, a more robust approach combining gel filtration, multi-angle light scattering, and SAXS (Fig. 4B and Fig. S4). The Kratky plot for apo *Bf*NFeoAB (Fig. 4C) exhibits a bell-shaped curve indicating a well-folded protein, with a width that suggests an elongated, non-globular conformation. In addition, the plot does not converge back to the q axis at high q values, possibly indicating some flexibility within the protein. Both analyses agree well with our Robetta models of apo *Bf*NFeoAB, in which *Bf*FeoA is folded separately and disparately of *Bf*NFeoB, and in which the *Bf*FeoA domain appears to sample different conformations dependent on the model. We surmise that this conformational flexibility is a result of the flexibility of the linker region that connects *Bf*FeoA and *Bf*NFeoB.

We then sought to determine the approximate size and molecular envelope of apo *Bf*NFeoAB. Using GNOM from the ATSAS package (45), the best solution for the protein maximal dimension (D_max_) of *Bf*NFeoAB was ≈80 Å, similar to the longest dimension observed of our elongated Robetta models. GASBOR (46) was then subsequently used to generate 10 independent *ab initio* envelopes (Fig. 4D, gray mesh). The overall shape of the envelope is consistent with an elongated shape, and we were able to identify easily a region in the envelope that strongly resembled the shape of the FeoA domain (Fig. 4D, overlay). To determine which of our Robetta models best fit the *ab initio* envelope, we used SUPCOMB (47), from the ATSAS package, to overlay each model with the envelope (Fig. 4D). In parallel, the online FoXS server (48,49) was used to determine the fit between the experimental scattering data and the theoretical scattering data calculated for each model. Our best Robetta model, which is shown in Fig. 4A, had a χ^2^ value of 1.81, indicating a good fit within the *ab initio* envelope.

To determine the effects of nucleotide on the overall structure of *Bf*NFeoAB, we also repeated our HT SAXS experiments in the presence of GMP-PNP and GDP (Fig. S3A). Though our samples were not as homogenous as our apo protein (Fig. S3B), we noted a remarked compaction in the overall structure (Fig. S3C), especially in the presence of GMP-PNP. These observations led us to hypothesize that FeoA could interact with NFeoB in a nucleotide-dependent manner and could lead to a compaction in structure, and we sought to test this hypothesis and to characterize the sites of this interaction.

### Hydrogen-Deuterium Exchange Mass Spectrometry

Given our observations that apo *Bf*NFeoAB exists as an elongated conformer in solution, and that the conformation of the construct appears to change in the presence of nucleotide, we next sought to map the nucleotide-dependent conformational changes and structural dynamics using hydrogen-deuterium exchange mass spectrometry (HDX-MS). To do so, we incubated apo protein with excess GDP or GMP-PNP (a non-hydrolyzable GTP analog) and compared the uptake of solvent deuterium of these forms of the protein to that of the apo form of the protein at different time points (10 s to 2 hr). After incubation, quenching, and digestion to obtain peptides, difference plots reveal the differential percent deuterium uptake in the apo protein compared to the GMP-PNP-bound form (Fig. S5) and the GDP-bound form (Fig. S6). Significant differences in deuterium uptake levels were then mapped onto our *Bf*NFeoAB model in a time-dependent manner for both the GMP-PNP- and GDP-bound forms, and these results reveal major and intriguing differences in the response of the protein based on nucleotide status and are described below.

Binding of the non-hydrolyzable GTP analog GMP-PNP elicits increases in protein protection over a slow time period and throughout the entire protein, suggestive of large changes in conformational dynamics that are consistent with protein compaction and our HT-SAXS data (Fig. S3). In the presence of GMP-PNP, minimal protection from deuterium uptake is observed over the 10 s to 1 min timeframe (Fig. 5A). Significant protection occurs in the GDI domain, in the region containing the additional α-helix and disordered loop (residues 270-323), and in part of the G4 motif (responsible for H-bonding with the guanine nucleotide), which is likely a result of nucleotide recognition. At the 10 min timepoint (Fig. 5B), increased protection is observed in the GDI domain. Moreover, regions of FeoA and the G-protein domain are also protected. These regions include four of the five residues comprising the hydrophobic clamp in FeoA (Ile^27^, Met^30^, Ile^59^, and Leu^61^), the G1 motif (responsible for binding to the α- and β-phosphate of GTP), the G3 motif (responsible for binding to the γ-phosphate of GTP and Mg^2+^), most of the Switch II region, the region between Switch II and the G4 motif, the PxxP motif (where we posit interactions with the hydrophobic clamp of FeoA occur), and the G5 motif (responsible for H-bonding to the guanine nucleotide). The protection observed at 10 min is consistent with nucleotide binding as most of the G-protein motifs exhibit protection. Interestingly, it is clear that protection of FeoA and the PxxP motif is observed on the same timescale, and these are the few regions that are not directly involved in nucleotide binding, suggesting protein-protein interactions occur in this region. Protection within the G2 motif (responsible for binding to the γ-phosphate of GTP and Mg^2+^) is observed only at the 1 hr and 2 hr timepoints (Fig. 5C). Under no conditions did we observe significant protection within the linker region or within the Switch I region in the presence of GMP-PMP. Notably, across all time points we only observe significant protection and no deprotection in GMP-PNP-bound *Bf*NFeoAB, indicating a compaction of structure in this form, consistent with our HT SAXS data.

**Fig. 5.**
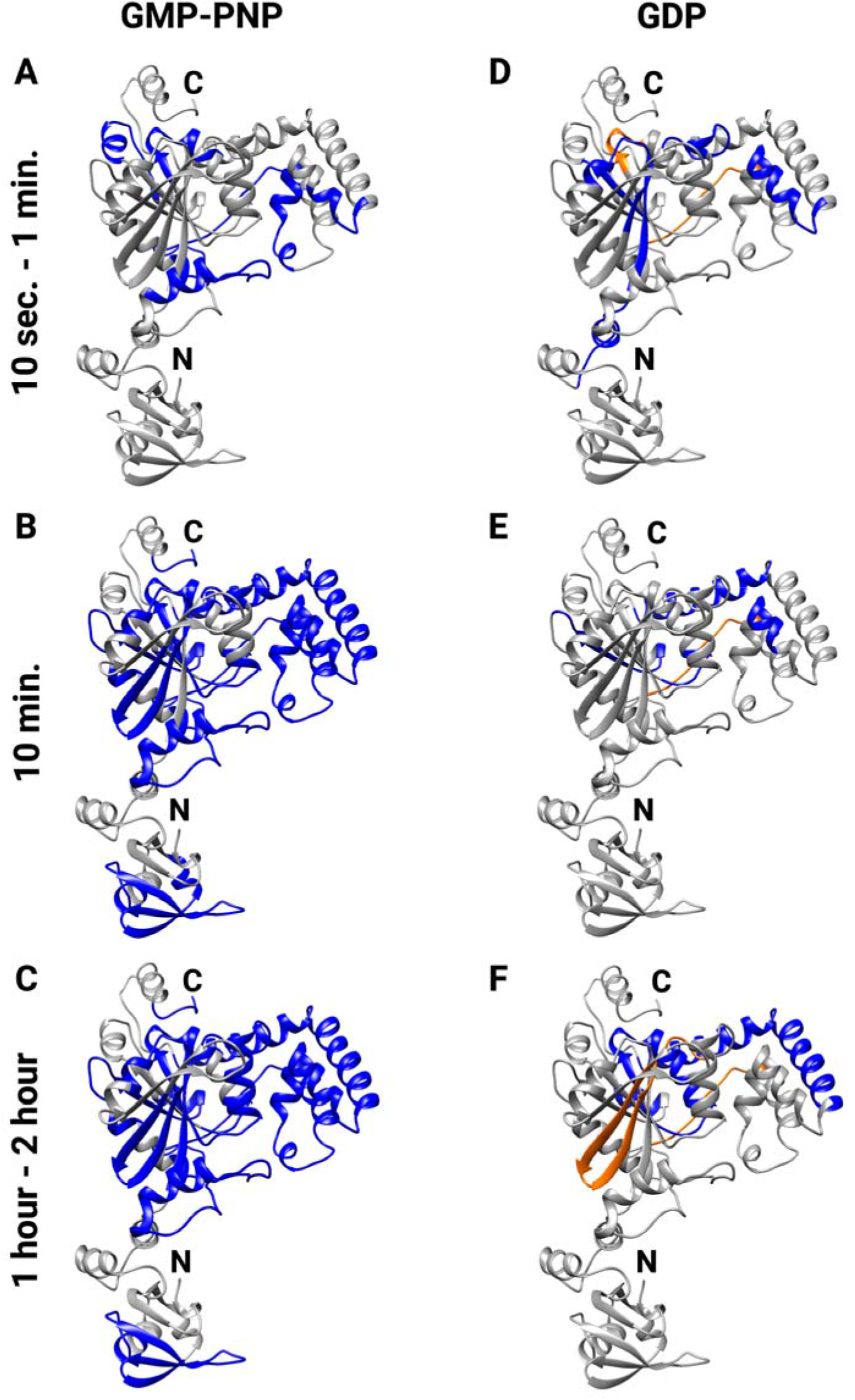
HDX-MS data on *Bf*NFeoAB indicate nucleotide- and time-dependent differences in protein protection (blue) and deprotection (orange), which are mapped onto the apo *Bf*NFeoAB Robetta model. **A**. Protection from deuterium uptake in the presence of GMP-PNP at both 10 s and 1 min. Most of the protection is observed in the GDI domain. **B**. Continued protection (compaction) is observed of *Bf*NFeoAB within 10 min, including the FeoA domain, the G-protein domain, and the GDI domain. **C**. After 1-2 hr, most of the GDI domain exhibits protection, as well as the Switch II region, and the key G-protein motifs. **D**. Deprotection of *Bf*NFeoAB is only observed in the presence of GDP, beginning as early as 10 s – 1 min, while minimal protection is observed. **E**. Within 10 min, deprotection is observed in the beginning of the GDI domain, while protection within in the rest of the protein is limited. **F**. After 1-2 hr, deprotection in the Switch I region is observed, as well as protection within the GDI domain. Labels ‘N’ and ‘C’ refer to the N- and C-termini, respectively.

In contrast, binding of GDP elicits a mixture of protein protection and deprotection and, importantly, does not engage FeoA. In the presence of GDP, we observed minimal protection within the GDI domain over the 10 s to 1 min timeframe (Fig. 5D), similar to the GMP-PNP bound form. However, we also noted protection within part of the linker region and the beginning of the G-protein domain, motifs G1, G3, and G4, and the beginning of the Switch II region. Unlike in the presence of GMP-PNP, we observed deprotection in the area adjacent to the Switch II region (residues 197-202) and in the random coil that feeds into the beginning of the GDI domain, consistent with our HT-SAXS data indicating the GDP-bound protein is more elongated than the GMP-PNP-bound protein (Fig. S3). This behavior could mimic the protein response post GTP hydrolysis, which would induce conformational changes for GDP release. At 10 min (Fig. 5E), protection is only observed in the region between Switch II and the G4 motif (residues 196-218), and within the PxxP motif. Lastly, by the 1 hr and 2 hr timepoints (Fig. 5F) deprotection is observed between the G2 and G3 motifs (residues 148-161), while protection is observed in the G3 motif, the beginning of the Switch II region, between Switch II and the G4 motif, between the G4 motif and the PxxP motif, and in the GDI domain. No protection or deprotection of FeoA is observed at any time point, suggesting FeoA does not interact with NFeoB in the GDP-bound state, which could poise the protein for GDP dissociation and/or nucleotide exchange.

### Rate of GTP hydrolysis

Finally, as no FeoA-FeoB fusion has been enzymatically characterized, we sought to examine the rate of GTP hydrolysis of *Bf*NFeoAB to garner insight into function. Using 1D-^31^P NMR spectroscopy, we performed a full, continuous kinetic characterization of *Bf*NFeoAB (Fig. 6). In this assay, signals indicative of GTP (−5.98, -10.91, and -19.39 ppm; corresponding to γ-, α-, and β-phosphates, respectively) slowly reduced in intensity as signals indicative of GDP (−6.08 and -9.92 ppm; corresponding to β- and α-phosphates, respectively) and inorganic phosphate (P_i_) (1.91 ppm) increased at the same slow rate. Due to its clear and distinct position, the P_i_ signal was integrated and plotted with respect to time in order to assess the rate of hydrolysis. Consistent with findings reported previously by Lau and coworkers (19), no auto-hydrolysis of GTP was observed under these conditions, and *Bf*NFeoAB exhibited remarkably slow GTP hydrolysis (*k*_*cat*_^*GTP*^ (0.71 ± 0.09) ×10^−3^ s^-1^) (Fig. 6, Table 1). As some NFeoBs are not strictly GTPases but, instead, are proposed NTPases (22,23), we also tested for ATPase activity but did not observe any protein-dependent ATP hydrolysis (data not shown). Interestingly, the rate of *Bf*NFeoAB-catalyzed GTP hydrolysis is at least an order of magnitude slower than all other observed NFeoBs, with the exception of *Kp*NFeoB (21,22,24). However, *Kp*NFeoB, a clear outlier, is known to bind its cognate FeoC (50) and we have shown that intact *Kp*FeoB can hydrolyze GTP *ca*. 100 ×10^−3^ s^-1^ (27). Moreover, many previous studies have reported increased GTPase activity of NFeoB upon replacement of NaCl with KCl in the reaction mixture (19,51), and this K^+^-dependent activation was attributed to two conserved Asn resides (51) that are also found in the sequence of *Bf*NFeoAB (Fig. S2). Therefore, we sought to test whether K^+^ stimulation could alter the rate of *Bf*NFeoAB-catalyzed GTP hydrolysis. To determine the influence of K^+^ on activity of *Bf*NFeoAB, NMR assays were conducted under identical conditions to those previously employed (*vide supra*) except that NaCl was replaced by KCl (Fig. S7). The rate of hydrolysis in K^+^ and Na^+^ were identical, demonstrating that *Bf*NFeoAB GTP hydrolysis is not activated in the presence of K^+^, like many NFeoBs. Thus, *Bf*NFeoAB appears to hydrolyze GTP slower than most NFeoBs, and access to the key Asn residue responsible for potassium stimulation is likely blocked from solvent. Given our findings that the binding of GTP analogs induces interactions between FeoA and NFeoB, we hypothesize that the presence of FeoA reduces the rate of GTP hydrolysis by protein-protein interactions that result in either direct or indirect occlusion of the nucleotide triphosphate.

**Table 1.**
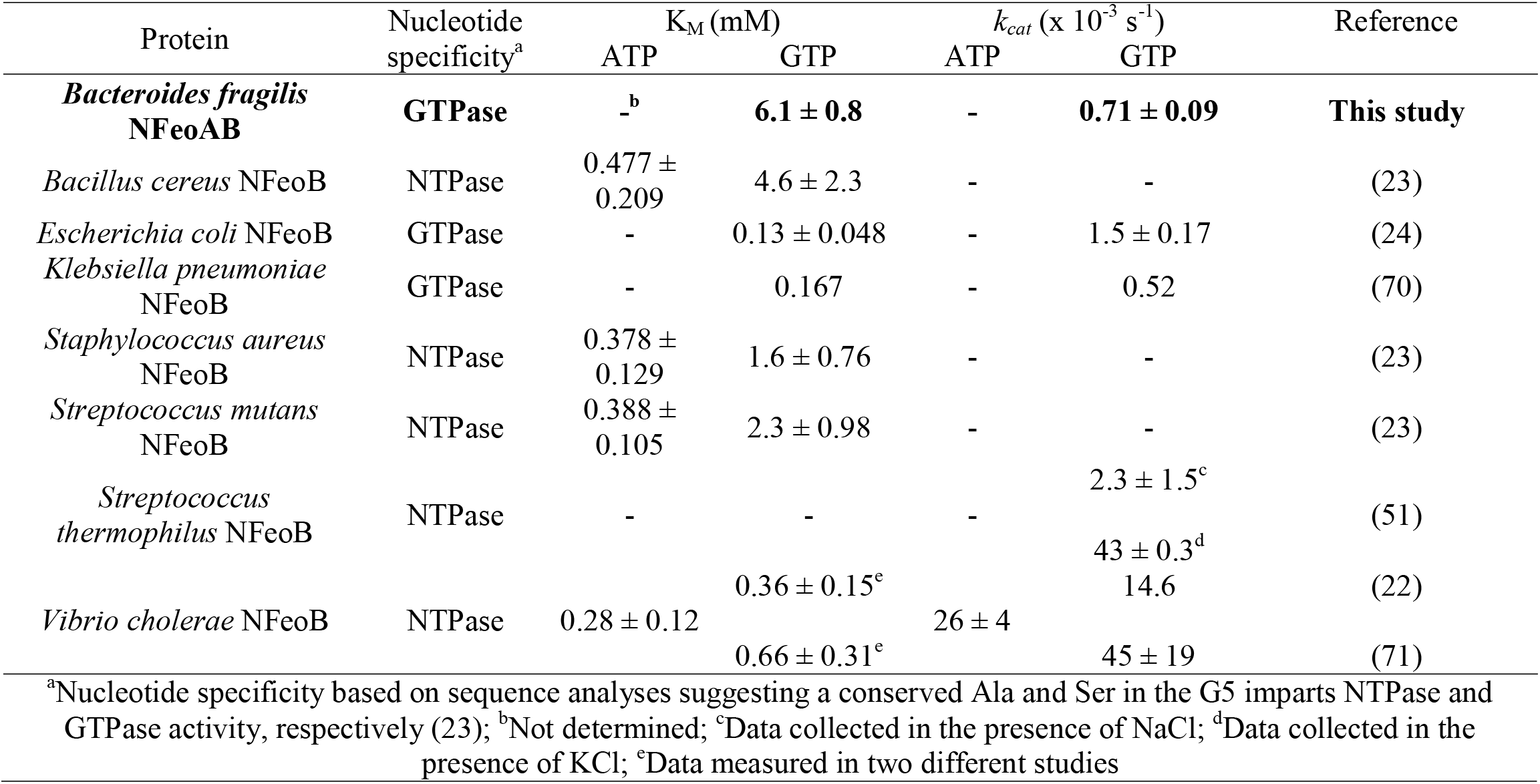
A comparison of the nucleotide-hydrolyzing enzymatic parameters of NFeoB proteins to those of *Bf*NFeoAB.

**Fig. 6.**
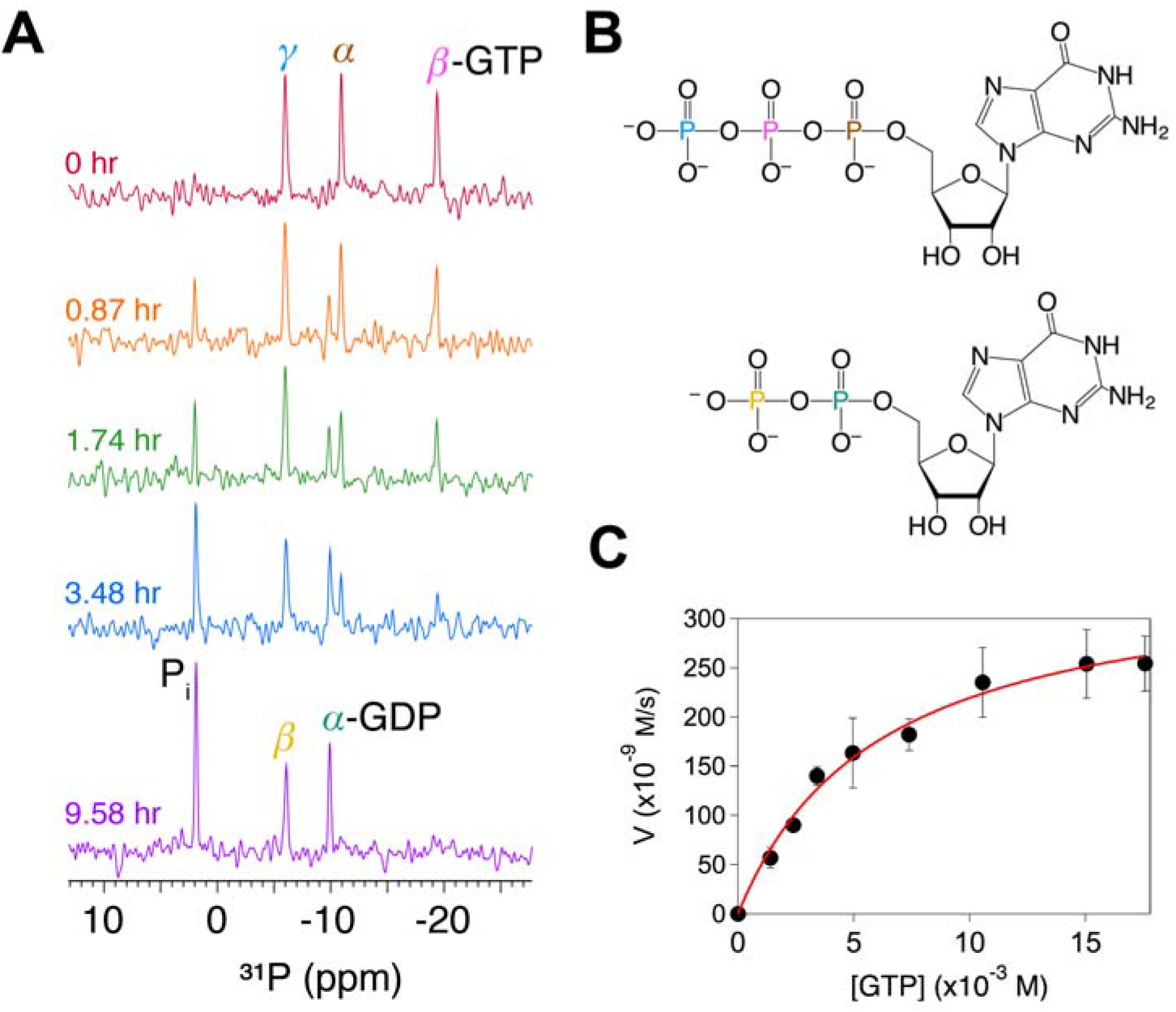
Characterization of the GTPase activity of *Bf*NFeoAB by nuclear magnetic resonance (NMR) spectroscopy indicates that nucleotide hydrolysis is exceedingly and notably slow. **A**. Exemplar of the 1D-^31^P NMR data used to determine the kinetics of 500 μM *Bf*NFeoAB in the presence of 5 mM GTP at 310 K. There is progressive reduction in the intensity of α-, β-, and γ-GTP phosphate chemical shifts (brown, magenta, and blue, respectively) until GTP is completely hydrolyzed to GDP (α and β in teal and yellow, respectively) and inorganic phosphate (P_i_; black). **B**. Structures of GTP and GDP color-coded to correspond to the phosphate chemical shifts in panel **A. C**. The Michaelis-Menten profile of *Bf*NFeoAB-catalyzed GTP hydrolysis, which demonstrates a maximum velocity (*V*_*max*_) of 0.35 ± 0.02 μM/s and a Michaelis constant (K_M_) of 6.1 ± 0.8 mM.

## DISCUSSION

The function of FeoA vis-à-vis FeoB has garnered considerable debate and yet failed to reach a consensus. Given its conserved SH3-like fold (19,20,42), which typically mediates protein-protein interactions in other systems, our lab and others have speculated that FeoA may interact with FeoB to alter function. In support of this notion, *in vivo* studies have demonstrated the ability of FeoA to interact with FeoB in stand-alone tripartite systems (52-54). However, when probed at the *in vitro* level, at least two studies have examined the role of FeoA and its effect on FeoB-catalyzed GTP hydrolysis in stand-alone tripartite systems (19,22), but virtually no effect was noted. It is possible that, in these tripartite systems, the lack of a membrane and/or the inability to reconstitute the correct multi-component protein system may have shrouded FeoA function. To circumvent this problem, we sought to probe the role of FeoA by taking advantage of a system that exists as a naturally occurring fusion.

We used multiple approaches to characterize a naturally occurring FeoA-FeoB fusion for the first time, with our first focus on structural determination. To decide initially which fusion protein to target, we undertook a bioinformatics approach, which revealed that these fusions are more widespread in bacteria than previously thought, and that the FeoA-FeoB fusion from *Bacteroides fragilis* would be a good representative target. After cloning, expression, and purification, this protein was used for crystallization, SEC-SAXS, HDX-MS, and enzymatic assays. As there has been much debate over the oligomeric state of NFeoB/FeoB (whether monomer or trimer), (27,40,41,50,54-57) it is worth noting that in our hands, *Bf*NFeoAB exists predominantly as a monomer (calculated MW 48.4 kDa ± 0.7 kDa) in solution at both high and low protein concentrations, though a small amount (estimated <10 %) exists as a trimer that is not in dynamic equilibrium with its monomeric form. Nevertheless, we tried to crystallize both, but despite exhaustive crystallization trials we were only able to crystallize the FeoA domain of this protein. The overall structure of *Bf*FeoA (Fig. 3) is similar to other FeoA proteins that have been determined by NMR or X-ray crystallography (19,20), including the hydrophobic cleft putatively involved in protein-protein interactions, suggesting that the FeoA domain in an FeoAB fusion may function similarly to stand-alone FeoA proteins.

In the absence of a crystal structure of the intact *Bf*NFeoAB, we turned to Robetta to generate homology models of our protein (Fig. 4A). In these models, FeoA appears to sample several different conformations, and this flexibility likely explains our difficulty in crystallizing the intact protein. However, this flexibility is likely linked to function, and we hypothesized interactions of FeoA and NFeoB could be nucleotide mediated. To investigate nucleotide-mediated conformational changes in *Bf*NFeoAB, we utilized SEC-SAXS and HDX-MS. Our SEC-SAXS data allowed us to determine the overall low-resolution structure of apo *Bf*NFeoAB in solution (Fig. 4), which matches our best-fit Robetta model, both of which indicate the FeoA and NFeoB do not interact in the absence of nucleotide. However, HT-SAXS data suggested a nucleotide-mediated interaction, which we probed with HDX-MS.

Our HDX-MS experiments clearly indicated nucleotide-mediated changes in *Bf*NFeoAB that are distinct based on whether nucleotide is intact (GMP-PNP) or hydrolyzed (GDP) and provide insight into the solution behavior of the fusion protein in response to nucleotide. In both the presence of GMP-PNP and GDP, protection occurs as early as 10 s and 1 min, predominantly in the GDI domain. We posit that this could transduce a signal to the TM region of FeoB and such a signal could “turn on” Fe^2+^ transport (Fig. 7). It is only in the presence of GMP-PNP that we observe protection within FeoA, most of the G-protein motifs, and the PxxP epitope. This behavior indicates that GMP-PNP (a proxy for GTP) binding induces structural and dynamic changes that, together with our SAXS data showing GMP-PNP-dependent compaction, indicate an FeoA-NFeoB interaction. In contrast, the effect of GDP on protection and deprotection in *Bf*NFeoAB is not nearly as dramatic. Similar to the binding of GMP-PNP, the binding of GDP initiates protection of the GDI domain that we posit that this transduces a signal to the TM region of FeoB to “turn off” Fe^2+^ transport (Fig. 7). The deprotection observed in the Switch I region in the presence of GDP could also be involved in the release of hydrolyzed nucleotide. Our data does suggest dynamic behavior in the Switch regions, which could have downstream effects on both nucleotide binding and release, consistent with several NFeoB crystal structures (57-59).

**Fig. 7.**
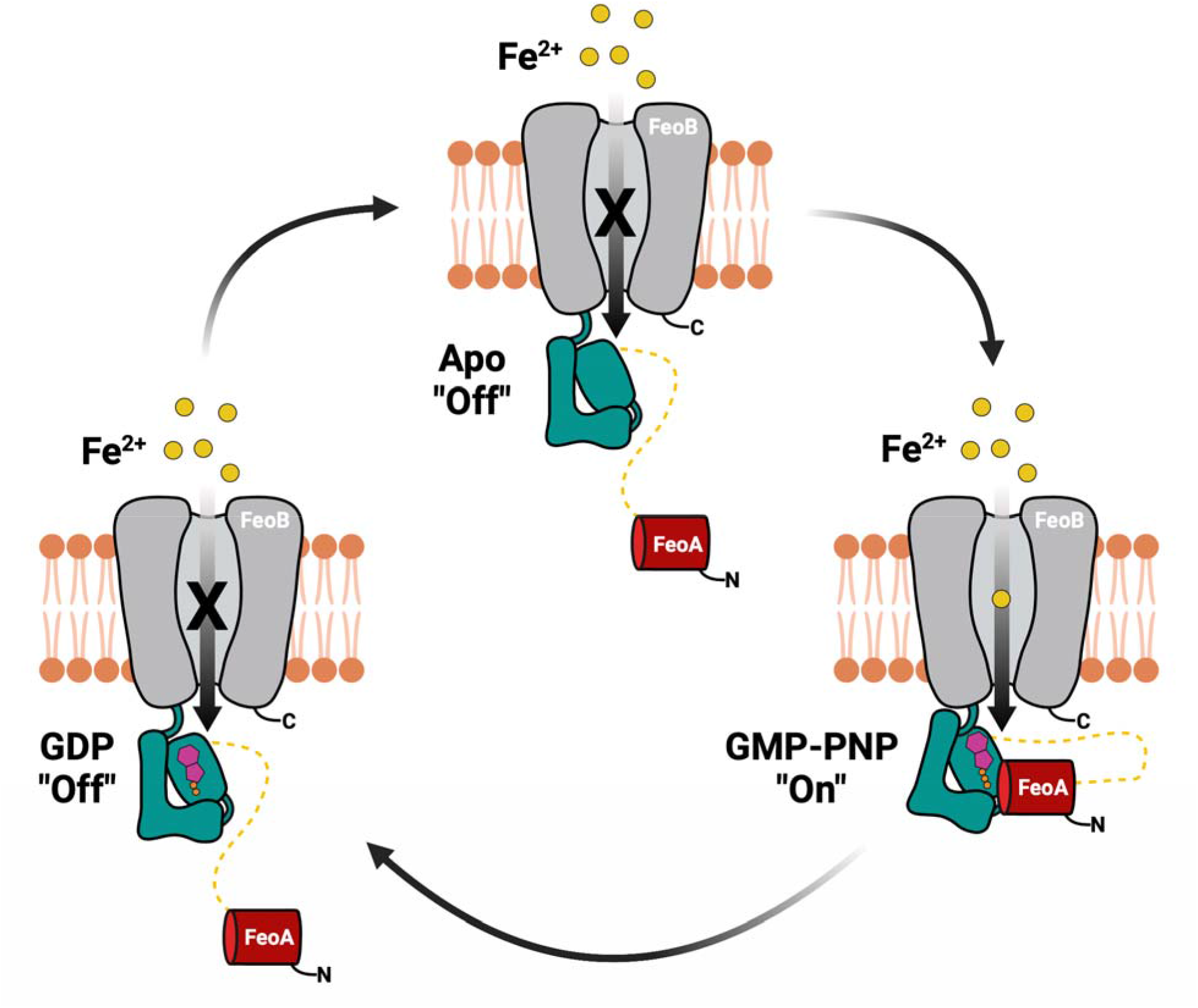
A model of the proposed FeoA-based regulatory function on iron transport of *Bf*FeoAB. In the absence of nucleotide (apo; top), FeoA (red) does not interact with NFeoB (teal), thus transport of Fe^2+^ (yellow) through the transmembrane domain (gray) is in an “off” state. Binding of GMP-PNP (magenta and orange; right), a GTP-analog, to NFeoB induces protein-protein interactions between NFeoB and FeoA. We posit that this GMP-PNP-bound state sends a signal to the transmembrane region to “turn on” Fe^2+^ transport. GDP binding to NFeoB (magenta and orange; left) transduces a signal the transmembrane domain to “turn off” Fe^2+^ transport before dissociation of FeoA-NFeoAB interactions, loss of GDP, and a return of the transporter to the apo state.

In combination with our biophysical analyses and our enzymatic data, we believe these results have strong implications for the function of FeoA. The rate of GTPase activity of *Bf*NFeoAB and its affinity to GTP (assuming K_M_ ∼ K_d_) are significantly lower than that of other NFeoBs (Table 1). We believe this phenomenon is explained by the role of FeoA rather than the lack of additional stimulatory factors. First, this and other fusion proteins do not contain an additional ORF in their operons encoding for FeoC, unlike the tripartite Feo systems. Second, neither the use of other nucleotides nor the presence of K^+^ elicits increased hydrolysis rates comparable to other NFeoBs. Third, our HDX-MS data show significant protection of FeoA and the PxxP motif of NFeoB, in addition to expected regions within the nucleotide binding site, indicating operative protein-protein interactions are occurring. Taken together, these findings support our hypothesis that FeoA may regulate the rate of GTP hydrolysis via interaction with NFeoB in a nucleotide-dependent manner, which likely controls iron transport across the membrane (Fig. 7). In particular, FeoA may function to slow GTP hydrolysis in order to keep the transporter “turned on” in a manner similar to eukaryotic GPCRs. After sufficient intracellular iron is then accumulated, an unknown downstream signal could dislodge FeoA from NFeoB to then “turn off” the transporter (Fig. 7). If operative, these protein-protein interactions could be targeted for therapeutic developments to treat infections. For example, multiple organisms like *B. fragilis* are commensal and colonize the human gut, but if they enter an environment outside of the gastrointestinal tract, they can cause bacteremia, which is becoming increasingly resistant to antibiotic treatments (39). The targeting of FeoA-NFeoB interactions could represent one way to treat these infections by starving the bacterium of this source of iron. However, future work on the intact membrane protein is necessary to probe this hypothesis further.

## EXPERIMENTAL PROCEDURES

### Materials

The codon-optimized gene encoding for the N-terminal soluble domain of *Bacteroides fragilis* FeoAB (*Bf*NFeoAB; Uniprot identifier A0A0K6BRR9) was commercially synthesized by GenScript. The pET-21a(+) expression plasmid was purchased from EMD-MilliporeSigma. A C43 (DE3) *E. coli* expression cell line carrying a deletion of *acrB*, an endogenous *E. coli* multidrug exporter, (C43 (DE3) Δ*acrB*) was provided by Prof. Edward Yu (Case Western Reserve University). Ampicillin and β-D-l-thiogalactopyranoside (IPTG) were purchased from RPI and used as received. Additional materials for cellular growth, protein expression, and protein purification were purchased from MilliporeSigma, VWR and/or Fischer Scientific and used as received. Sparse-matrix crystallization screens were purchased from Hampton, stored at 4°C, and allowed to equilibrate at room temperature before use. Guanosine triphosphate (GTP), guanosine diphosphate (GDP), and 5’guanylyl-imidodiphosphate (GMP-PNP) were purchased from MilliporeSigma, stored at -20°C and used as received. D_2_O was purchased from MilliporeSigma and used as received.

### Bioinformatics

FeoA proteins existing as naturally occurring fusions were identified by searching the InterPro Database (accessed February 2021) for all domain architectures containing the FeoA protein (InterPro ID: IPR007167) fused to the G-protein domain of FeoB (InterPro ID: IPR030389). FeoA-like proteins fused to DtxR-type helix-turn-helix domains (InterPro ID: IPR022687), iron dependent repressor, metal binding and dimerization domains (InterPro ID: IPR001367) and/or other non-Feo-related proteins were manually removed from the data. The resulting FeoA-containing domain architectures were downloaded from the InterPro database and analyzed. Hierarchical taxonomic information was extracted directly from the truncated database. Multiple sequence alignments were performed using ClustalW (60) and visualized in the JalView suite (v. 2.11.1.4, (61)). To determine portions of the FeoAB sequences that belong to cytosolic or membrane domains, the transmembrane helix prediction server (TMHMM, v. 2.0) (62) was used.

### Cloning, expression, and purification of BfNFeoAB

The presence of the FeoA protein fused to the N-terminal soluble domain of *Bacteroides fragilis* (*Bf*NFeoAB) was identified from the full-length FeoAB protein sequence (Uniprot identifier A0A0K6BRR9). Using TMHMM, v. 2.0 (62), amino acid residues 1-438 were determined to comprise the extent of the cytosolic, soluble fusion domain. The codon-optimized gene encoding for these residues plus an engineered DNA sequence encoding for a C-terminal tobacco etch virus (TEV)-protease cleavage site (ENLYFQS) was synthesized by GenScript. This gene was then subcloned into the ampicillin-resistant, IPTG-inducible pET-21a(+) expression plasmid using the NdeI and XhoI restriction sites, encoding for a C-terminal (His)_6_-affinity tag used for affinity purification. This expression plasmid bearing *Bf*NFeoAB was subsequently transformed into chemically-competent C43 (DE3) Δ*acrB E. coli* expression cells, plated on Luria-Bertani (LB) agar plates containing ampicillin (final concentration of 100 μg/mL) and incubated overnight at 37°C.

Single colonies obtained from these plates were then used to inoculate 100 mL of LB media containing ampicillin (final concentration of 100 μg/mL). This preculture was grown overnight at 37°C with shaking of 200 RPM and subsequently used to inoculate 12 baffled flasks each containing 1 L of sterile LB media and supplemented with sterile ampicillin (final concentration of 100 μg/mL). These large-scale cultures were grown at 37°C with shaking of 200 RPM until the OD_600_ was ≈0.6-0.8. Cultures were then cold shocked for 2 hours at 4°C, and protein expression was induced by the addition of sterile IPTG to a final concentration of 1 mM. Flasks were then incubated overnight at 18°C with shaking of 200 RPM. Cells were harvested after ≈16-18 hours by centrifugation at 4800×*g* for 10 minutes at 4°C. Cell pellets were resuspended in resuspension buffer (50 mM Tris, pH 7.5, 300 mM NaCl, and 5% (v/v) glycerol), then flash-frozen in N_2(l)_ and stored at -80°C.

To initiate the purification of *Bf*NFeoAB, cells were thawed, homogenized, and solid phenylmethylsulfonyl fluoride (PMSF; 50-100 mg) was added prior to cell sonication at 4°C on a Q700 ultrasonic cell disruptor (QSonica) at an amplitude of 80%, 30 sec pulse on, 30 sec pulse off for 12 minutes. The cellular lysate was then clarified by ultracentrifugation at 163000×*g*, 4°C for 1 hour and the supernatant was applied to a 5 mL immobilized metal affinity chromatography (IMAC) HisTrap HP column (Cytiva) charged with Ni^2+^ and pre-equilibrated at 4°C with 5 column volumes (CVs) of wash buffer (50 mM Tris, pH 8.0, 300 mM NaCl, 5% (v/v) glycerol, 1 mM TCEP HCl) with an additional 21 mM imidazole. The column was then washed with an additional 8 CVs of wash buffer with an additional 21 mM imidazole followed by a wash with 16 CVs of wash buffer with an additional 50 mM imidazole. Protein was eluted from the column with elution buffer (50 mM Tris, pH 8.0, 300 mM NaCl, 5% (v/v) glycerol, 1 mM TCEP, 150 mM imidazole). Fractions containing eluted *Bf*NFeoAB were concentrated at 4°C using a 15 mL Amicon 3 kDa molecular-weight cutoff (MWCO) spin concentrator (Millipore). Concentrated protein was then further purified by size exclusion chromatography (SEC) on a 120 mL Superdex 200 column (Cytiva) equilibrated with SEC buffer (25 mM Tris, pH 7.5, 300 mM NaCl, 2% (v/v) glycerol, 1 mM TCEP) operating at 4°C. Fractions containing monomeric *Bf*NFeoAB were then concentrated at 4°C using a 4 mL Amicon 3 kDa MWCO spin concentrator. Protein concentration and purity were determined using the Lowry assay and 15% sodium dodecyl sulfate polyacrylamide gel electrophoresis (SDS-PAGE) analysis.

### Crystallization, data reduction, and structural determination

SEC-purified *Bf*NFeoAB was concentrated to ≈20 mg/mL and screened for crystallization at room temperature using the vapor diffusion method in 96-well sitting drop trays using commercially available crystallization screens. Initial crystals were obtained in 0.2 M di-potassium phosphate and 2.2 M ammonium sulfate. These crystals were optimized by grid screening in 24-well sitting drop trays at room temperature using the vapor diffusion method. Medium-sized rectangular, crystals appeared in several wells after ≈11 months. Crystals were harvested, cryoprotected for ≈1 sec in a drop containing 1 M ammonium sulfate, 0.1 M di-potassium phosphate and 25% (v/v) glycerol, and frozen on N_2(l)_. Diffraction data were collected at the Advanced Photon Source (APS), Argonne National laboratory on LS-CAT beamline 21-ID-G. Data were automatically processed using Xia2 (63). Phasing was achieved by molecular replacement (MR) using Phenix Phaser (64) with *Clostridium thermocellum* FeoA (PDB ID: 2K5L) as an input search model. After an initial MR solution was identified, further model building was accomplished using Phenix AutoBuild (64). Iterative rounds of manual model building and refinement were accomplished in Coot (65) and Phenix Refine (64), respectively, until model convergence. The final model consists of residues 1-74 of the FeoA portion of *Bf*NFeoAB. This structure has been deposited in the Protein Data Bank (PDB ID 7R7B). Data collection and refinement statistics are provided in SI Table 1.

### Homology modeling

The structural prediction of apo *Bf*NFeoAB was determined using comparative modeling, a method used for targets with homologs in the PDB, via the Robetta online server (43,44). Five structures of *Bf*NFeoAB were generated using comparative modeling, each with high confidence. Each model was tested for its agreement to the experimentally-determined structure of *Bf*FeoA (*vide supra*) and for its fit into the *ab initio* generated molecular envelope that was created from SEC-coupled small angle X-ray scattering data (*vide infra*).

### Small-angle X-ray scattering

High throughput (HT-) and SEC-coupled small-angle X-ray scattering (SEC-SAXS) data were collected at the Advanced Light Source (ALS), Lawrence Berkeley National Laboratory, on the SIBYLS beamline 12.3.1. A suite of samples each containing 60 μL of *Bf*NFeoAB at concentrations ranging from 4-6 mg/mL were screened after passage along a PROTEIN KW-803 column equilibrated with SAXS buffer (25 mM Tris, pH 7.5, 150 mM NaCl, 2% (v/v) glycerol, 1 mM TCEP HCl) using an autosampler. Eluent was split 2:1 between the X-ray synchrotron radiation source (SAXS) and a series of four inline analytical instruments: 1) Agilent 1260 series multiple wavelength detector (MWD); 2) Wyatt Dawn Helos multi-angel light scattering (MALS) detector; 3) Wyatt DynaPro Titan quasi-elastic light scatterings (QELS) detector; and 4) Wyatt Optilab rEX refractometer. Samples were examined with λ = 1.03 Å incident light at a sample-to-detector distance of 1.5 m resulting in scattering vectors, q, ranging from 0.01 Å^-1^ to 0.5 Å^-1^ where the scattering vector is defined as q=4πsinθ/λ and 2θ is the measured scattering angle. Data were collected in 3 s exposures over the course of 40 min. SEC-SAXS chromatograms were generated and initial SAXS curves were analyzed using SCÅTTER (66,67). Additionally, UV, MALS, QELS, and differential refractive index data was collected and analyzed.

Scattering curves were analyzed using SCÅTTER (66,67) and GNOM (45) to generate Guinier and Kratky plots and to determine the radius of gyration (*R*_*g*_) and the maximum particle dimension (*D*_*max*_). *Ab initio* molecular envelopes were generated using GASBOR (46) and averaged with DAMAVER (68) from the ATSAS package and were displayed using Mac PyMOL (v. 2.4.1) and UCSF Chimera. *Bf*NFeoAB Robetta models were overlayed with SUPCOMB (47), also part of the ATSAS package and displayed using Mac PyMOL (v. 2.4.1). The online Fast SAXS Profile Computation with Debye Formula (FoXS) (48,49) server was used to determine which Robetta model (*vide supra*) best fit the generated *ab initio* envelopes.

### Hydrogen deuterium exchange coupled to mass spectrometry (HDX-MS)

To begin, undeuterated controls were performed for peptide identification to obtain a sequence coverage map for *Bf*NFeoAB. The experimental workflow is as follows: 2 μL of 20 μM *Bf*NFeoAB in 25 mM Tris pH 7.5, 300 mM NaCl, 2% (v/v) glycerol, 1 mM TCEP was diluted with 98 μL of ice-cold quench (100 mM glycine pH 2.5, 1 M guanidine-HCl, 5mM TCEP). After 1 min, the 100 μL dilution was injected into a Waters HDX nanoACQUITY UPLC (Waters, Milford, MA) with in-line protease XIII/pepsin digestion (NovoBioAssays LLC). Peptic fragments were trapped on an ACQUITY UPLC BEH C18 peptide trap and separated on an ACQUITY UPLC BEH C18 column. A 7 min, 5% to 35% acetonitrile in 0.1% formic acid gradient was used to elute the peptides directly into a Waters Synapt G2-Si mass spectrometer (Waters, Milford, MA). MS^e^ data were acquired with a 20 to 30 V ramp trap collision energy (CE) for high energy acquisition of product ions and continuous lock mass (Leucine-Enkephalin) for correction of mass accuracy. Peptides were identified using the ProteinLynx Global Server 3.0.3 (Waters). A filter of 0.3 fragments per residue was applied for peptide processing in the DynamX 3.0 software (Waters).

Hydrogen-deuterium exchange reactions for apo *Bf*NFeoAB and the protein in complex with GMP-PNP were performed by manual injections. The same reactions of the apo protein and the protein in complex with GDP were acquired with a LEAP autosampler controlled by the Chronos software. The reaction workflow for both manual and autosampler injections was as follows: 4 μL of 10 μM protein in complex with 5 mM GMP-PNP or 5 mM GDP was incubated in 36 μL of 25 mM Tris in D_2_O (99.99%), pD 7.5, 300 mM NaCl, 2% (v/v) glycerol and 1 mM TCEP. The 40 μL reaction was quenched at various times with 60 μL of 100 mM glycine pH 2.5, 2.5 M guanidine-HCl and 5mM TCEP. All the deuteration reactions were carried out at 25°C at five reaction time points (10 s, 1 min, 10 min, 1 hr, and 2 hr). Following quenching of the deuterated samples, the 100 μL quenched reaction was injected and LC/MS acquisition was performed in the same manner as the undeuterated controls. The five deuteration time points were acquired in triplicate. Fully deuterated controls were performed for normalization purposes. The normalized percent deuterium uptake (%D) for each peptide, at incubation time *t*, was calculated as described in the equation below:

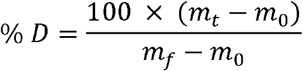

where *m*_*t*_, *m*_0_ and *m*_*f*_ are the centroid masses at incubation time *t*, the undeuterated control, and the fully deuterated control, respectively. The reaction workflow for the fully deuterated controls was as follows: 10 μL of 60 μM *Bf*NFeoAB was incubated with 10 μL of 25 mM Tris pH 7.5, 7.84 M guanidine-HCl and the protein was incubated overnight. Subsequently, 4 μL of the unfolding reaction was diluted with 36 μL D_2_O buffer, pD 7.4, and allowed to deuterate for more than 2 hours. The reaction was quenched with 60 μL quench buffer and injected, with LC/MS acquisition performed as described above. The DynamX 3.0 software was used for spectral curation, centroid calculation, and deuterium uptake analysis of all identified peptides.

### GTPase assays

Samples for nuclear magnetic resonance (NMR) experiments were prepared in 100 mM Tris, pH 7.5, 300 mM NaCl, 100 mM MgSO_4_, 2% (v/v) glycerol, 1 mM TCEP, and 10% D_2_O in a 3 mm NMR tube. Experiments in which the contribution of K^+^ was monitored were carried out under the same conditions as outlined above except NaCl was replaced by KCl. The GTPase and ATPase activities of ≈500 μM *Bf*NFeoAB were monitored by 1D-^31^P NMR spectroscopy at 37°C using a 500 MHz Bruker DMX spectrometer equipped with a room temperature probe. NMR spectra were collected using 256 scans with a 10-30 minute delay between acquisitions, and data were processed using DataChord Spectrum Analyst (69). The velocity profiles were based on linear initial rates; to determine Michaelis-Menten kinetics, GTP concentrations were varied from 1.5 mM to 18 mM. Over these substrate concentrations, initial velocity measurements were plotted vs. substrate concentration (GTP or ATP) and fitted to the following equation:

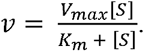

## Supporting information

Supplemental Data for Manuscript

## Acknowledgements

This work was supported by NIH-NIDCR grant R21 DE027803, NIH-NIGMS grant R35 GM133497, in part by NIH-NIGMS grant T32 GM066706 (A. E. S. and J. O. O), and by the University of Maryland Baltimore, School of Pharmacy Mass Spectrometry Center (SOP1841-IQB2014). This research used resources of the Advanced Photon Source, a U.S. Department of Energy (DOE) Office of Science User Facility operated for the DOE Office of Science by Argonne National Laboratory under Contract No. DE-AC02-06CH11357. Use of the LS-CAT Sector 21 was supported by the Michigan Economic Development Corporation and the Michigan Technology Tri-Corridor (Grant 085P1000817). SAXS experiments were conducted at the Advanced Light Source (ALS), a national user facility operated by Lawrence Berkeley National Laboratory on behalf of the Department of Energy, Office of Basic Energy Sciences, through the Integrated Diffraction Analysis Technologies (IDAT) program, supported by the DOE Office of Biological and Environmental Research. Additional support comes from the National Institute of Health project ALS-ENABLE (P30 GM124169) and a High-End. Instrumentation Grant S10OD018483. Sequence searches utilized both database and analysis functions of the Universal Protein Resource (UniProt) Knowledgebase and Reference Clusters (http://www.uniprot.org) and the National Center for Biotechnology Information (http://www.ncbi.nlm.nih.gov/). NMR experiments were carried out at the University of Maryland Baltimore County Molecular Characterization and Analysis Complex.

## Supporting Information

Supporting information is available online, including:

*Bf*FeoA data collection and refinement statistics (Table S1)

The 2F_o_-F_c_ electron density map of *Bf*FeoA (Fig. S1)

Partial multiple sequence alignments (MSAs) of the soluble, N-terminal domains of non-fused

FeoB proteins and select FeoAB fusion proteins (Fig. S2)

Experimental high-throughput (HT) SAXS data for apo and nucleotide-bound *Bf*NFeoAB (Fig. S3)

SEC-R_*g*_ profile for apo *Bf*NFeoAB (Fig. S4)

Difference plots of apo *Bf*NFeoAB percent deuterium uptake (%D_apo_) minus GMP-PNP-bound

*Bf*NFeoAB percent deuterium uptake (%D_GMP-PNP_) (Fig. S5)

Difference plots of apo

*Bf*NFeoAB percent deuterium uptake (%D_apo_) minus GDP-bound *Bf*NFeoAB percent deuterium uptake (%D_GDP_) (Fig. S6)

*Bf*NFeoAB GTP hydrolysis in the presence of K^+^ and Na^+^ (Fig. S7)

## Conflict of interest

The authors declare no competing financial interests.

### Abbreviations

The abbreviations used are:

GDP: guanosine diphosphate
GMP-PNP: 5’guanylyl-imidodiphosphate
GTP: guanosine triphosphate
IMAC: immobilized metal affinity chromatography
IPTG: isopropyl β-D-l-thiogalactopyranoside
NFeoAB: soluble N-terminal GTP-binding domain of FeoB fused to FeoA
NFeoB: soluble N-terminal GTP-binding domain of FeoB
NMR: nuclear magnetic resonance
PMSF: phenylmethylsulfonyl fluoride
SAXS: small-angle X-ray scattering
SDS-PAGE: sodium dodecyl sulfate polyacrylamide gel electrophoresis
SEC: size-exclusion chromatography
SEC-SAXS: SEC-coupled small-angle X-ray scattering
R.M.S.D.: root-mean-square deviation
TEV: Tobacco Etch Virus
Tris: tris(hydroxymethyl)aminomethane
TCEP: tris(2-carboxyethyl)phosphine.

