## Supplemental Data for Manuscript for "Biochemical and structural characterization of the fused *Bacteroides fragilis* NFeoAB domain reveals a role for FeoA"

Running title: Characterization of *B. fragilis* NFeoAB

^*^To whom correspondence should be addressed: Aaron T. Smith: Department of Chemistry and Biochemistry, University of Maryland, Baltimore County, Baltimore, Maryland, 21250; Tel: 410-455-1985;

**Keywords:** Feo, iron, protein-protein interactions, SAXS, SH3, iron transport

| **Table 1** | |
| --- | --- |
| **Data Collection and Refinement Statistics for *Bf*FeoA (1-74)** | |
| **Data Collection** |  |
| Beamline | APS 21-ID-G |
| Wavelength (Å) | 0.97856 |
| Space group | *C*_121_ |
| Cell Dimensions |  |
| *a*, *b*, *c* (Å) | 92.58, 29.55, 67.43 |
| α, β, γ (˚) | 90, 128.89, 90 |
| Resolution (Å) | 46.18-1.50 (1.53-1.50) |
| *R*_merge_ | 0.122 (1.185) |
| *I/σ(I)* | 8.1 (1.5) |
| Completeness (%) | 99.7 (97.8) |
| **Refinement** |  |
| Resolution (Å) | 36.03-1.50 (1.55-1.50) |
| No. reflections | 23,079 |
| *R*_work_ | 0.192 |
| *R*_free_ | 0.236 |
| No. atoms |  |
| Protein | 1,214 |
| Glycerol | 12 |
| Phosphate | 10 |
| Water | 154 |
| Average B-factors (Å^2^) | 24.00 |
| R.m.s. deviations |  |
| Bond lengths (Å) | 0.006 |
| Bond angles (˚) | 0.878 |
| Ramachandran plot |  |
| Favored | 72 (97.22%) |
| Allowed | 2 (2.78%) |
| Outlier | 0 (0%) |

**Table S1.** *Bf*FeoA data collection and refinement statistics.

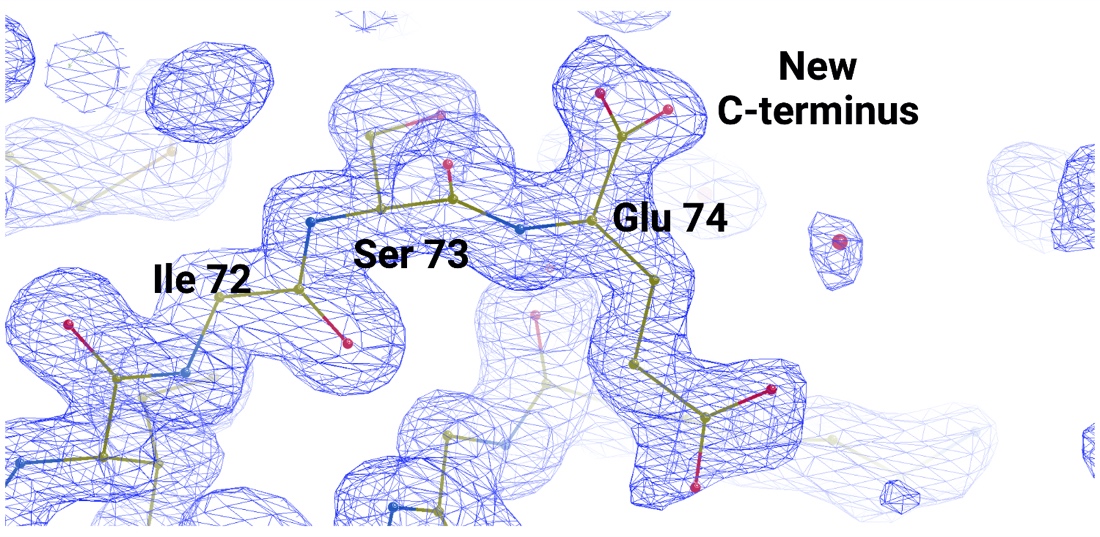

**Fig. S1.** The 2F_o_-F_c_ electron density map (blue) of *Bf*FeoA contoured to 1σ indicates the location of proteolytic cleavage. Glu 74 (shown here for chain B, but consistent throughout the ASU) forms the C-terminus of the polypeptide after proteolytic cleavage between residues 74 and 75 in the crystallization drop. Figure created with Biorender.

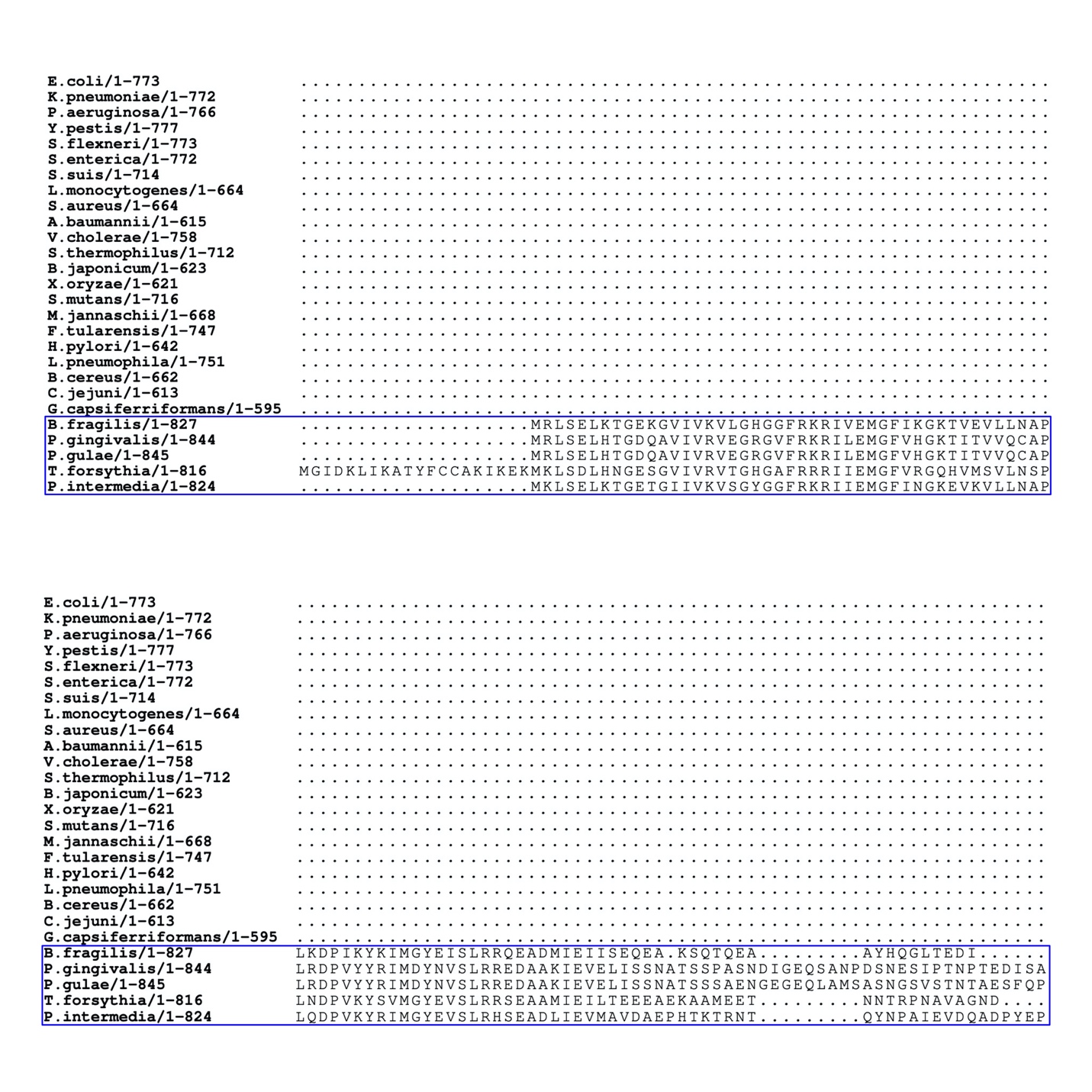

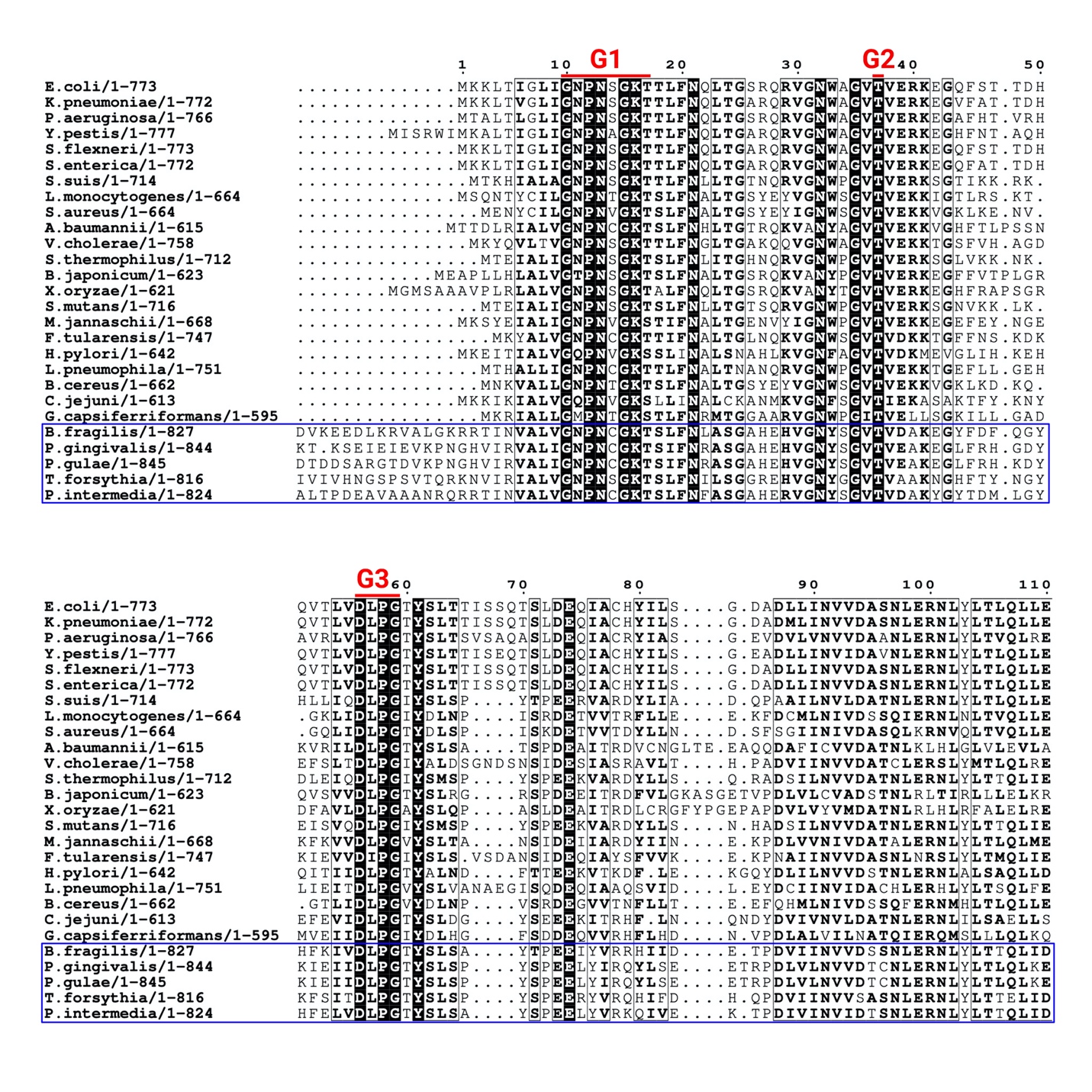

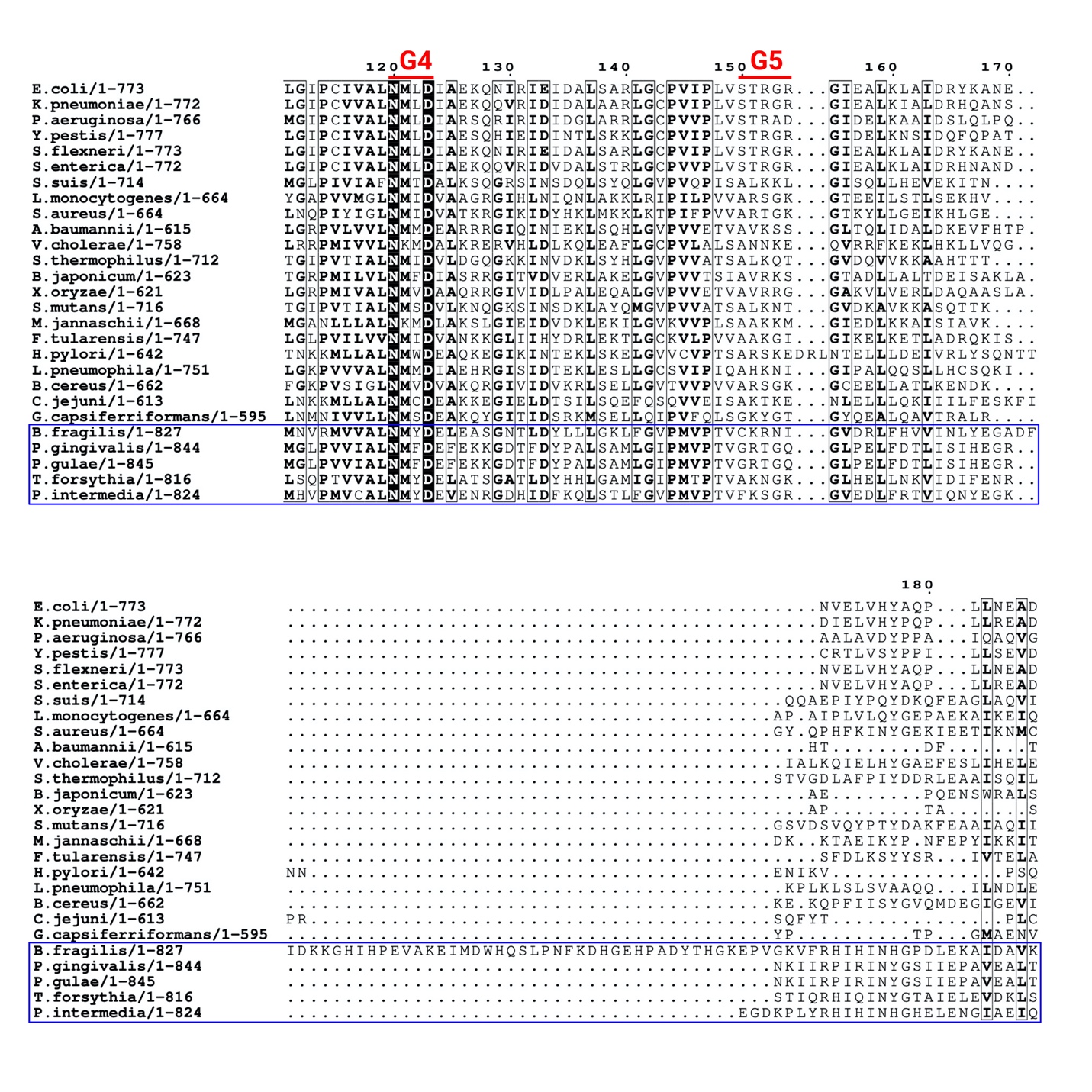

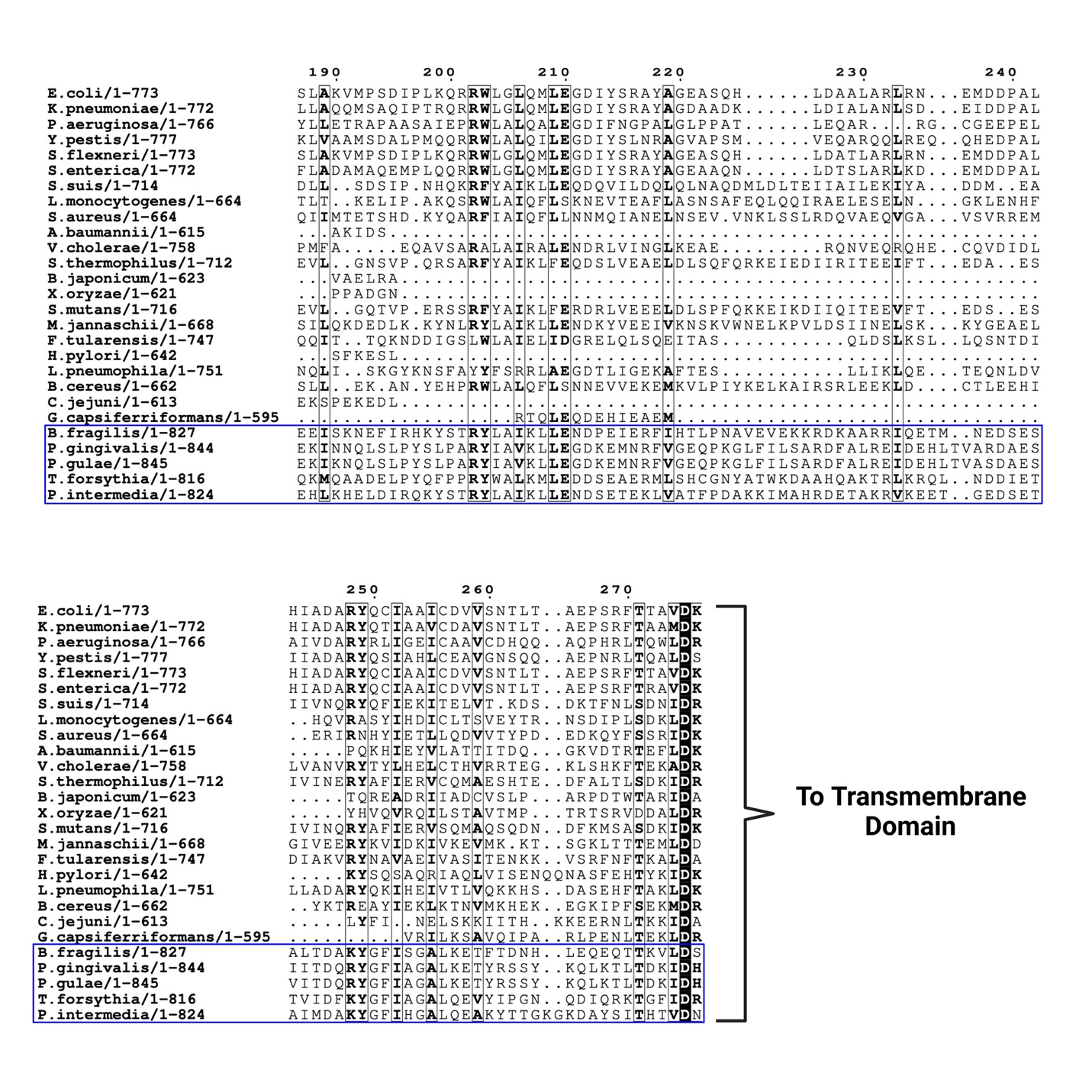

**Fig. S2.** Partial multiple sequence alignments (MSAs) of the soluble, N-terminal domains of non-fused FeoB proteins and select FeoAB fusion proteins (blue box). The top numbering is relative to *E. coli* FeoB (Uniprot ID P33650). The G motifs (G1-G5) are highlighted above the aligned sequences in red. MSAs were generated using ClustalW (1) and visualized in the JalView Suite (2) and the ESPript 3.0 online server (3). Figure created with Biorender.

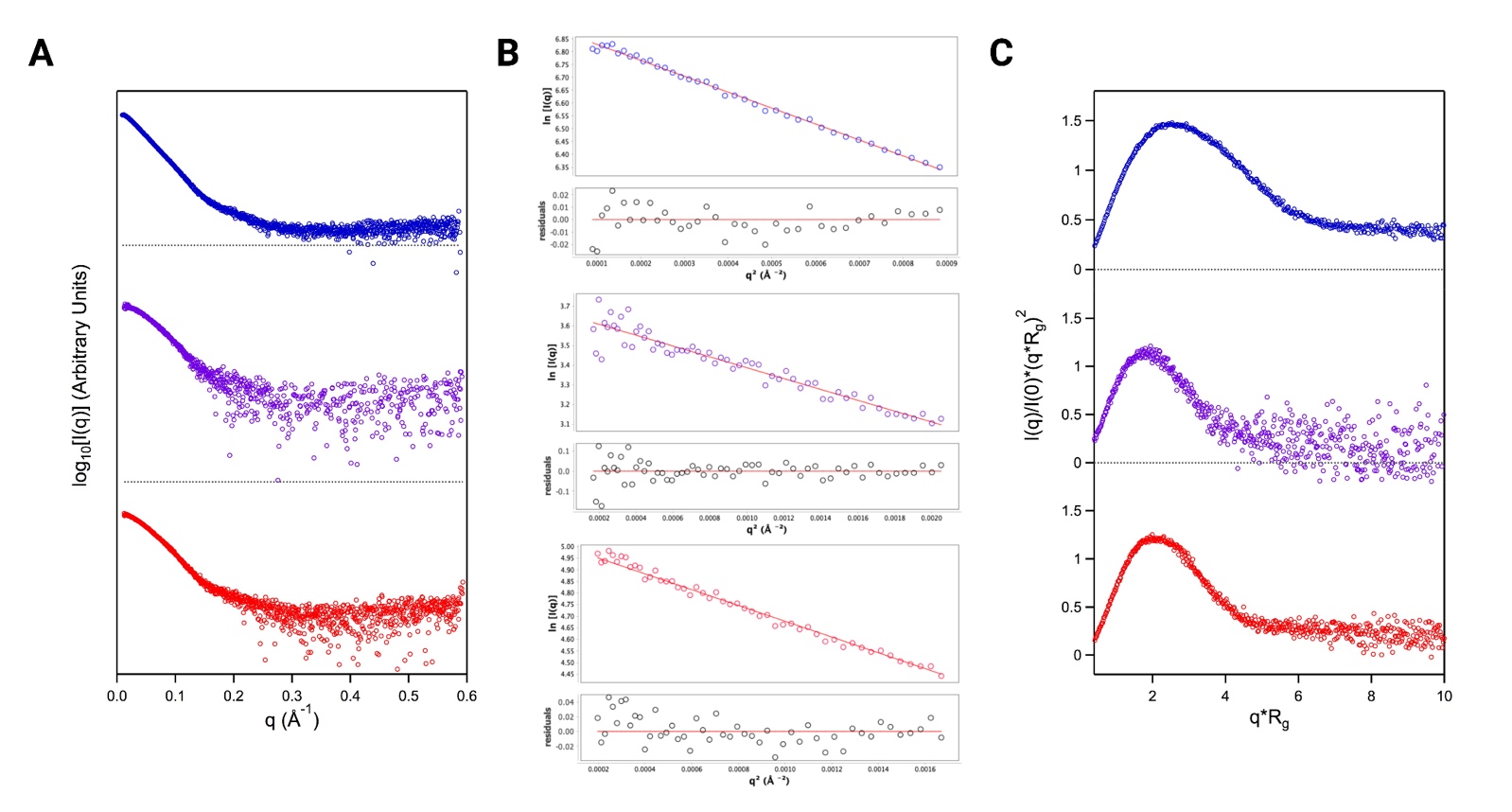

**Fig. S3.** Experimental high-throughput (HT) SAXS data for apo- and nucleotide-bound *Bf*NFeoAB. (**A**) log_10_ plots derived from experimental SAXS data of apo *Bf*NFeoAB (top, blue), GMP-PNP- (middle, purple), and GDP-bound (bottom, red) *Bf*NFeoAB. (**B**) Guinier fittings (solid red lines) and their respective fitted residuals (black circles) of apo *Bf*NFeoAB (top, blue), GMP-PNP- (middle, purple), and GDP-bound (bottom, red) *Bf*NFeoAB demonstrating negligible aggregation in our HT-SAXS samples. (**C**) The normalized Kratky plot of apo *Bf*NFeoAB (top, blue) suggests an elongated, flexible conformation while the normalized Kratky plots of GMP-PNP- (middle, purple) and GDP-bound (bottom, red) *Bf*NFeoAB suggest more compact, less flexible structures.

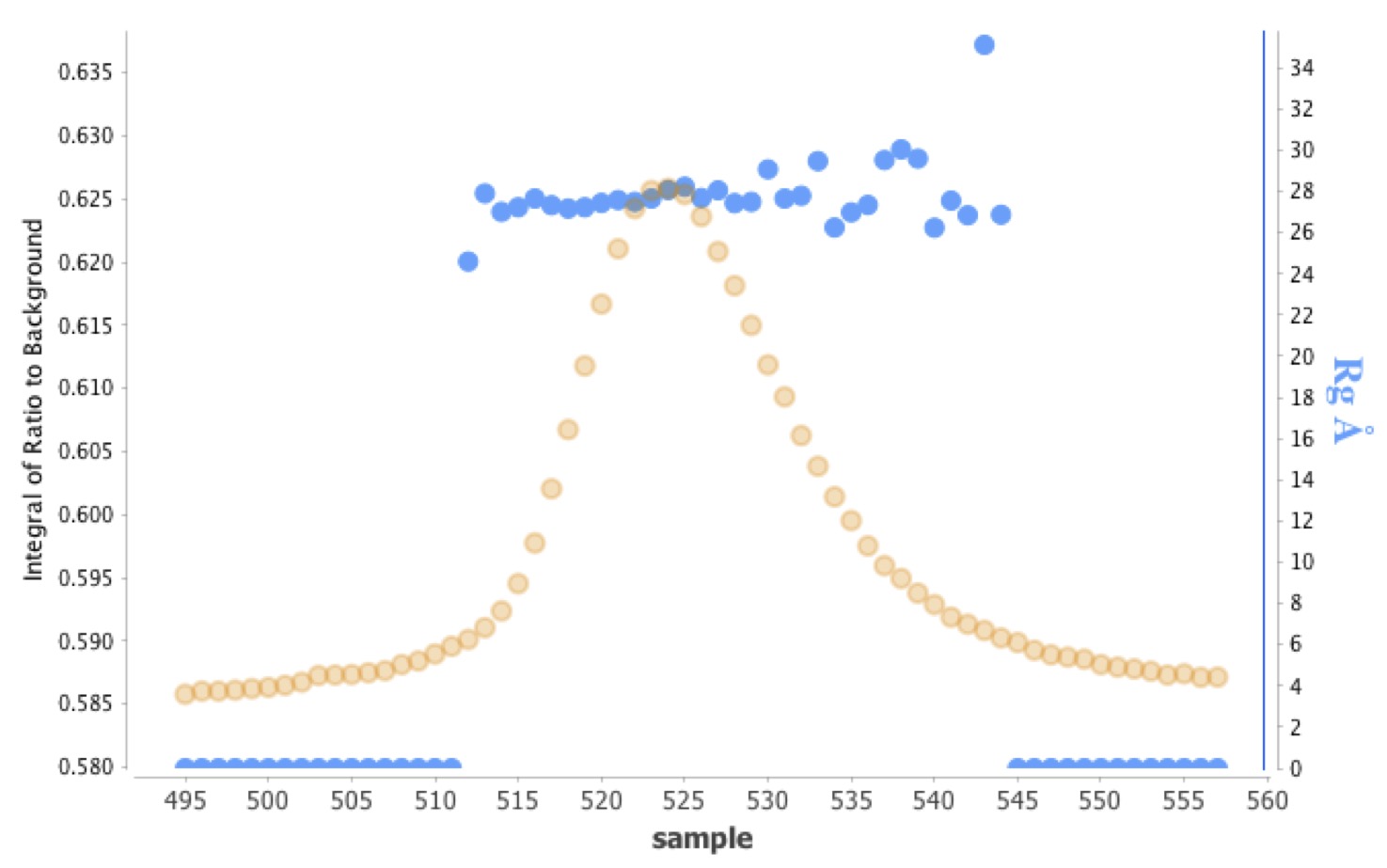

**Fig. S4.** SEC-R*_g_* profile for apo *Bf*NFeoAB. The SEC profile for apo *Bf*NFeoAB (orange circles) shows a homogenous, monodisperse sample. At the peak maximum, the R*_g_* values (blue circles) remain effectively constant at *ca*. 28 Å, further indicating homogeneity.

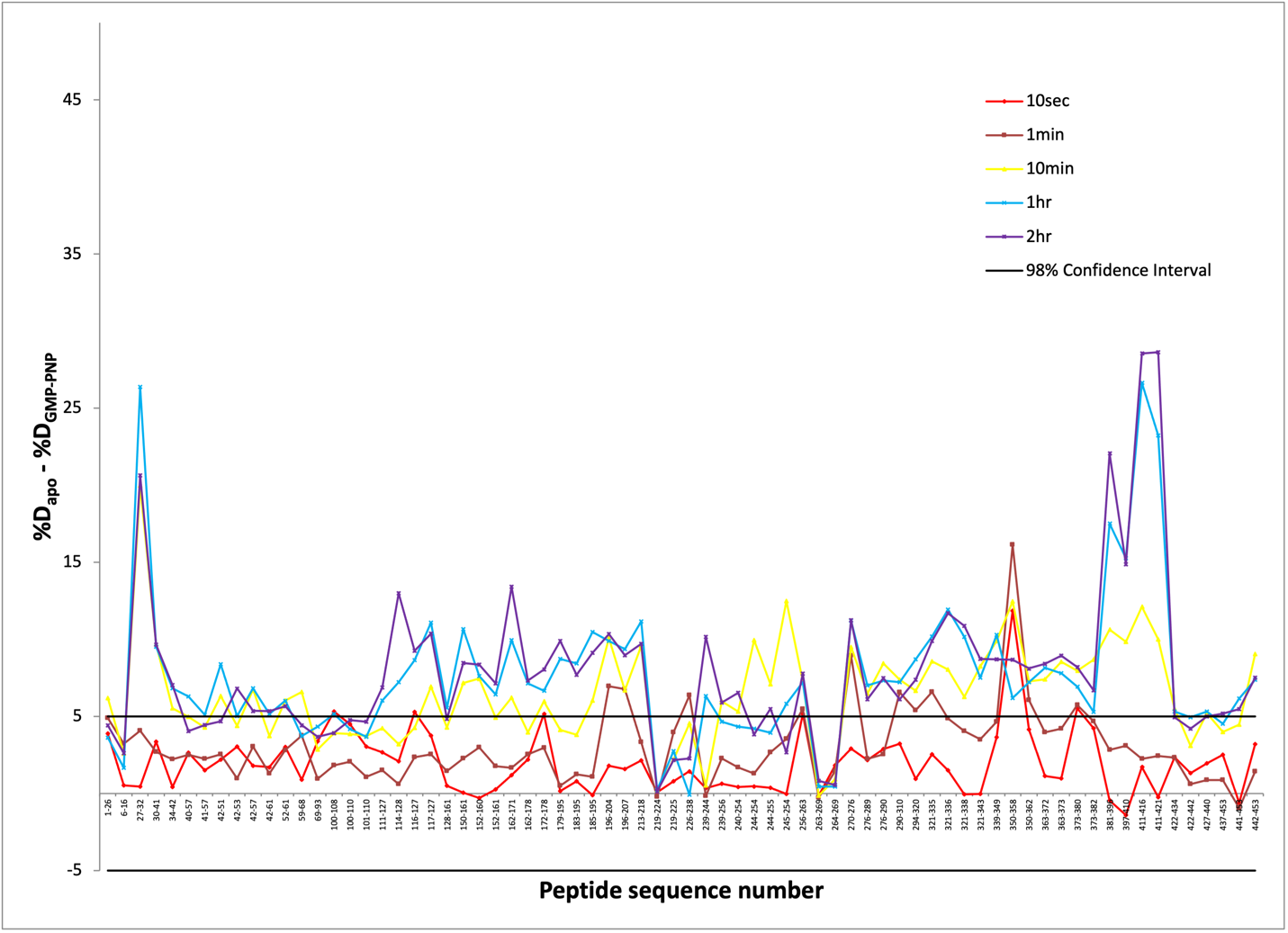

**Fig. S5.** Difference plots of apo *Bf*NFeoAB percent deuterium uptake (%D_apo_) minus GMP-PNP-bound *Bf*NFeoAB percent deuterium uptake (%D_GMP-PNP_). Individual peptic segments are plotted on the x-axis according to sequence number from N- to C- terminus. The time intervals represent the data after 10 s (red), 1 min (brown), 10 min (yellow), 1 hr (blue), and 2 hr (purple) of hydrogen-deuterium exchange reaction in each state. A 98% confidence interval (black line) was used for the determination of significance in the percent deuterium uptake difference.

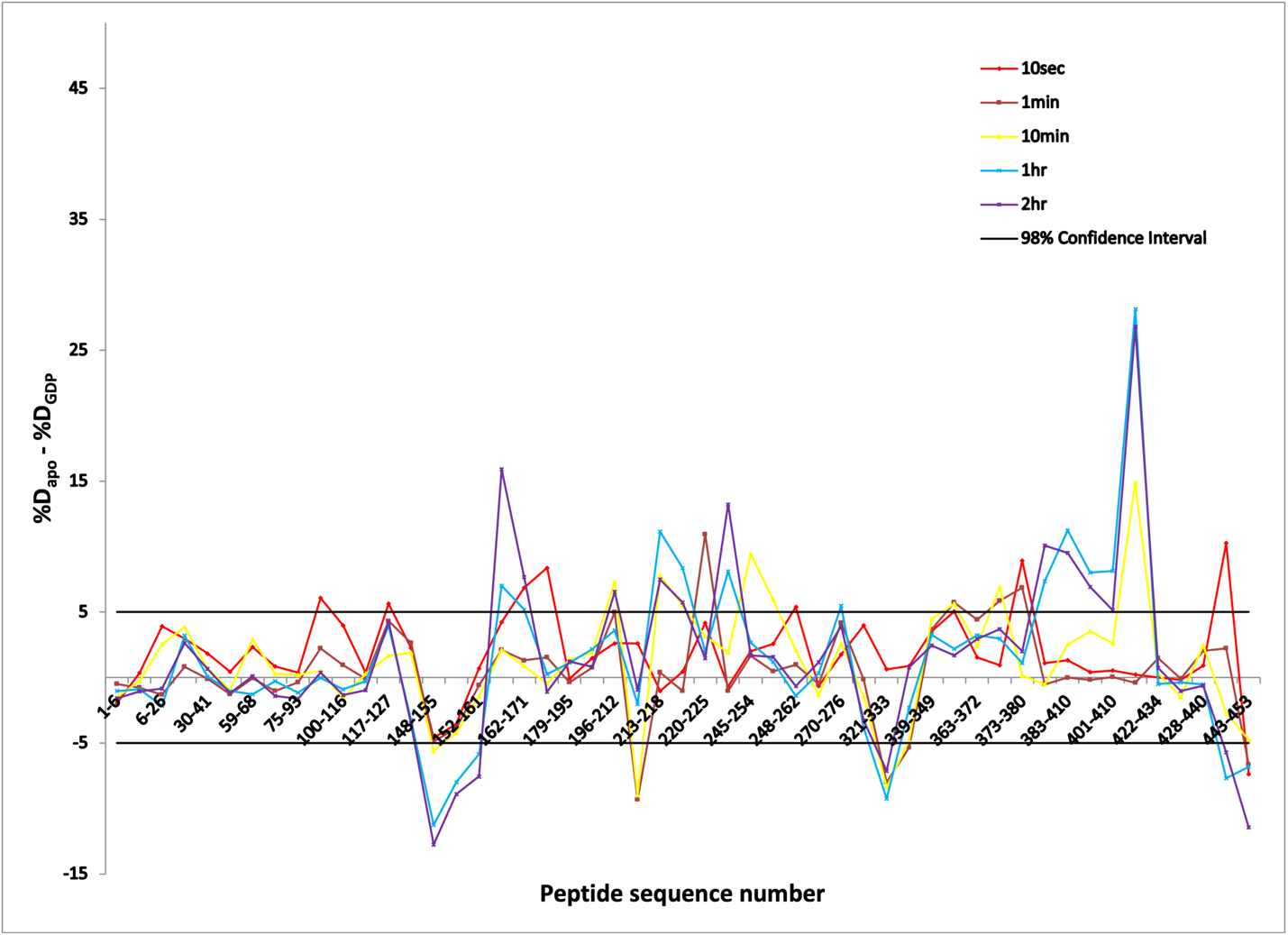

**Fig. S6.** Difference plots of apo *Bf*NFeoAB percent deuterium uptake (%D_apo_) minus GDP-bound *Bf*NFeoAB percent deuterium uptake (%D_GDP_). Individual peptic segments are plotted on the x-axis according to sequence number from N- to C- terminus. The time intervals represent the data after 10 s (red), 1 min (brown), 10 min (yellow), 1 hr (blue), and 2 hr (purple) of hydrogen-deuterium exchange reaction. A 98% confidence interval (black line) was used for the determination of significance in the percent deuterium uptake difference.

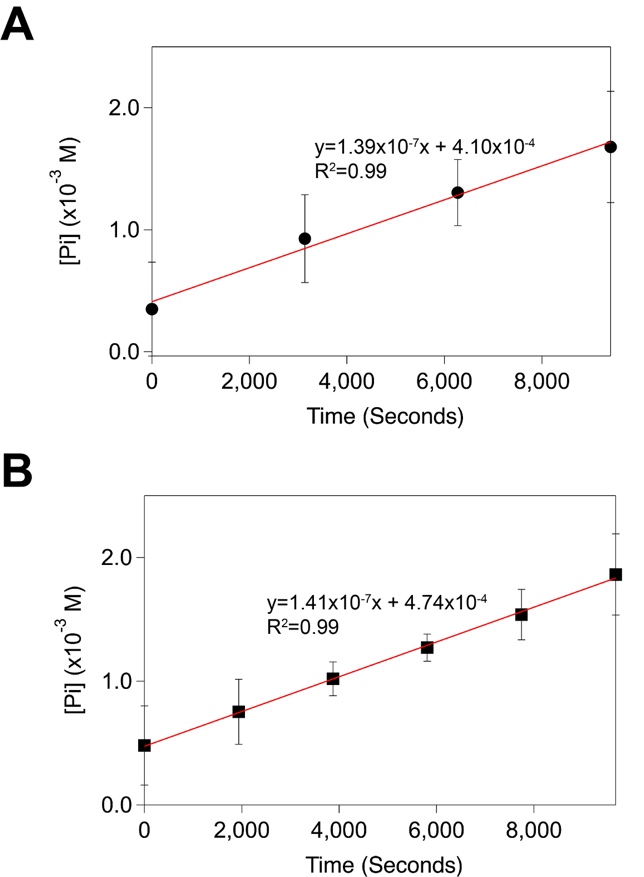

**Fig. S7.** The rate of *Bf*NFeoAB GTP hydrolysis is not increased by the presence of K^+^. (**A**) The reaction rate of *Bf*NFeoAB hydrolysis in the presence of 3.5 mM GTP was determined by plotting data collection time points against the area under the P_i_ signal in each 1D ^31^P-NMR spectrum. Sample conditions were: 100 mM Tris, pH 7.5, 300 mM NaCl, 100 mM MgSO_4_, 2% (v/v) glycerol, 1 mM TCEP, and 10% (v/v) D_2_O. Under these conditions, the rate of GTPase activity was linear and measured to be 0.14 ± 0.011 µM/s. (**B**) To determine the influence of K^+^ on GTP hydrolysis, the assay was carried out as described in panel **A** except that NaCl was replaced with KCl. The rate of GTP hydrolysis was measured to be 0.14 ± 0.003 µM/s, indicating that K^+^ does not affect *Bf*NFeoAB activity on GTP.
